# *Aedes albopictus* is rapidly invading its climatic niche in France: wider implications for biting nuisance and arbovirus control in Western Europe

**DOI:** 10.1101/2025.02.14.638223

**Authors:** Andrea Radici, Pachka Hammami, Arnaud Cannet, Grégory L’Ambert, Guillaume Lacour, Florence Fournet, Claire Garros, Hélène Guis, Didier Fontenille, Cyril Caminade

## Abstract

The Asian tiger mosquito, *Aedes albopictus,* is a competent vector of arboviruses, such as dengue. After its introduction into southern Europe, this invasive species has been rapidly spreading as well as causing autochthonous cases of arboviral diseases. Both *Ae. albopictus* presence and potential to transmit arboviruses are facilitated at warm temperatures, hence global warming is expected to affect their presence in temperate regions.

We use a climate- and environmental-driven mechanistic modelling framework to investigate the impact of recent climate change on *Ae. albopictus* range expansion and its potential to transmit dengue in Western Europe. We simulate climatic suitability, adult density and dengue transmission risk which we compare with a large ensemble of entomological and epidemiological observations over the past 20 years. Most importantly, we analyze a novel spatiotemporal dataset of colonized municipalities in metropolitan France to estimate the spread rate of *Ae. albopictus* and compare it with model predictions. Lastly, we analyze the sensitivity of entomological and epidemiological risk to changes in temperature, rainfall and human density.

Distribution of simulated mosquito populations and dengue transmission risk satisfactorily match entomological and dengue observations for Western Europe (AUC = 0.90 and 0.75 respectively). While lowlands in southern Europe were already climatically suitable for hosting *Ae. albopictus* around 2010, Western France, together with large populated cities, such as London, Zagreb and Vienna, have become suitable recently. Importantly, the accelerating colonization of *Ae. albopictus* in France may be approaching the limit of its theoretical climatic niche; future expansion will depend on the climate-driven enlargement of suitable areas. Area at risk of dengue transmission has recently expanded from the Mediterranean coasts over northern Spain and Western France. The sensitivity analysis suggests that climate change may expose medium-sized cities to the highest epidemiological risk; this finding is consistent with recently reported dengue outbreaks in Europe.

## Introduction

The Asian tiger mosquito *Aedes albopictus* is native to southeast Asian forests (Porretta et al., 2012). Over the past century, it has undergone genetic, phenotypic and behavioral adaptations, which favored its establishment outside its native range (Sherpa et al., 2019), in human habitats and temperate climates. Such features include laying eggs in man-made water reservoirs (Paupy et al., 2009) and producing diapausing eggs, which can withstand prolonged dry (Kraemer et al., 2019) or cold seasons (Batz et al., 2020; Medley et al., 2019). Facilitated by the worldwide shipping of used tires, *Ae. albopictus* successfully colonized areas in all continents except Antarctica (Kraemer et al., 2019; Sherpa et al., 2019).

Together with *Aedes aegypti*, *Ae. albopictus* is a competent vector of arboviral diseases, such as dengue, Zika and chikungunya (Bohers et al., 2024). Their ability to spread and adapt, coupled with their biting nuisance and competence to transmit arboviruses, makes the *Aedes* spp. the costliest among the invasive species (Diagne et al., 2021), with social and economic impacts expected to rise in the future (Roiz et al., 2024). Consequently, investigating and anticipating the spread of *Ae. albopictus* is a compelling issue in global public health (Ryan et al., 2019).

*Aedes albopictus* was likely introduced into Europe multiple times since its first establishment in Albania in 1979 (Sherpa et al., 2019). Following these introductions, *Ae. albopictus* spread like wildfire over southwestern Europe. Such spread was associated with occasional long-distance dispersal events, most likely facilitated by vehicular traffic (Roche et al., 2015; Roques & Bonnefon, 2016). The establishment of *Ae. albopictus* populations, combined with viral introductions by infected travelers returning from endemic countries, resulted in sporadic – but increasing in frequency and intensity – autochthonous cases of arboviruses (Angelini et al., 2007; Manica et al., 2017; Roiz et al., 2015). Worryingly, autochthonous cases of dengue have recently increased in Italy and France. 83 cases of dengue were reported over southern France in 2024 (Santé Publique France, 2024), associated with imported cases from the French Antilles, while 213 autochthonous dengue cases were reported in Italy in 2024, with about 199 cases occurring in Fano, in the Marches region (Sacco et al., 2024).

The life cycle of *Ae. albopictus* depends on a number of environmental factors (Cruz et al., 2024; Gizaw et al., 2024). The presence of water reservoirs, either rain-fed or man-fed, is indispensable for hosting the aquatic stages (Paupy et al., 2009). In temperate climates, the photoperiod regulates the production of diapausing eggs and their hatching (Lacour et al., 2015). As an ectotherm species, the life cycle of *Ae. albopictus* is primarily regulated by temperature (Mordecai et al., 2019). Being a species typical of tropical areas, climatic suitability for *Ae. albopictus* is favored at warm temperatures (Mordecai et al., 2017, 2019; Souza & Weaver, 2024). In addition, warm temperatures facilitate the transmission of arboviruses, since they reduce the extrinsic incubation period (EIP) in *Aedes* mosquitoes and accelerate their gonotrophic cycle (Mordecai et al., 2017).

Given the broad interest in anticipating public health risks induced by climate change (Souza & Weaver, 2024), there is a growing body of literature focusing on simulating the impact of a warming climate on *Ae. albopictus* (Barman et al., 2024; Caminade et al., 2012; Couper et al., 2021; Garrido Zornoza et al., 2024; Kraemer et al., 2019; Lamy et al., 2023; Metelmann et al., 2019) and its potential to transmit arboviruses at continental and regional scales (Blagrove et al., 2020; Caminade et al., 2017; Colón-González et al., 2021; Mordecai et al., 2017; Ryan et al., 2019; Zardini et al., 2024).

*Aedes albopictus* presence and abundance have been successfully modelled using both statistical (Kobayashi et al., 2002; Petrić et al., 2021; Ryan et al., 2019; Tran et al., 2020) and mechanistic (Brass et al., 2024; Da Re et al., 2022; Erguler et al., 2016; Metelmann et al., 2019; Tran et al., 2013) approaches. Some of these models have been used to investigate historical and future distribution and abundance of *Ae. albopictus* under different climate emission scenarios. Simulations usually show a general increase in suitable areas over the temperate fringes of the mosquito’s current distribution. For example, Metelmann et al. (2019) predicted that *Ae. albopictus* could theoretically spread to England in future, even if the species is not established there yet. Brass et al. (2024) suggests that recent climatic conditions are already suitable for *Ae. albopictus* in central Europe, while Zardini et al. (2024) suggest that central Europe may represent the northernmost limit for its presence.

Dengue transmission risk is usually assessed with the basic reproduction number R_0_. Some examples of integrating simulated mosquito density into R_0_ estimates have already been developed for dengue (Benkimoun et al., 2021). A multi-model multi-scenario inter-comparison study shows a northern shift of the dengue transmission belt in temperate Europe for the future (Colón-González et al., 2021). Another study estimates that, whatever the climate emission scenarios, transmission of arboviruses will be possible as far north as the Scandinavian peninsula by the 2050s (Ryan et al., 2019).

While scientific evidence suggests that future climatic conditions will become increasingly suitable for *Ae. albopictus* and the arboviruses it can transmit, the role of recent global warming in driving the current range expansion of *Ae. albopictus* is still under debate. *Aedes albopictus* showed remarkable adaptive capabilities, suggesting that its colonization process results from progressive adaptation of this species to new environments (Batz et al., 2020; Urbanski et al., 2012) rather than from the enlargement of already suitable areas. Despite the growing body of literature focusing on the impact of future climate change on the presence and epidemic activity of the Asian tiger mosquito, the contribution of recent climate change to its observed colonization of temperate zones has been largely overlooked.

In this study, we use a climate-driven mechanistic model (Metelmann et al., 2019) to assess the impact of recent climate change in determining the presence and the epidemic activity of *Ae. albopictus* in Western Europe. We first compare simulated mosquito distribution with presence/absence observations for Europe. In addition, we assess the capability of the vector model in reproducing observed seasonal and interannual variability using aggregated ovitrap data for different European locations. We employ a novel spatiotemporal dataset of colonized municipalities in metropolitan France to estimate the relative importance of recent climate change in driving the observed spread of *Ae. albopictus* at national scale. We then compare our R_0_ estimations for dengue with reported autochthonous cases of dengue in Europe. Finally, we conduct sensitivity experiments to disentangle the relative importance of climatic parameters and human population density on simulated suitability and epidemic risk. Based on our findings, we formulate recommendations for vector control operators and public health officers in France and Europe.

## Material and methods

### Entomological presence-absence data

Adult mosquito occurrence data was derived from the Global Biodiversity Information Facility (GBIF) repository up to 2024 (GBIF.org, 2024). GBIF collects species occurrence data from heterogeneous sources and does not provide absences. Therefore we used the European Centre for Disease Control (ECDC) data, collected and validated by experts of the VectorNet project (Wint et al., 2023), for defining absence of mosquitoes at the NUTS3 administrative level (ECDC, 2024b).

In metropolitan France, we employed a novel spatiotemporal dataset of colonized municipalities, from the national surveillance program, provided by the “*Direction générale de la santé”* (French Ministry of Health). This dataset includes a comprehensive list of municipalities and the corresponding year of colonization from 2004 to 2024. According to the national surveillance criteria, updated in the Decree of the 23^rd^ of July 2019, the Asian tiger mosquito is considered as established in a municipality if one of the following criteria is met: (i) eggs are collected in ovitraps during three consecutive inspections; (ii) larvae or adults are observed in traps in a 150 m radius around a positive ovitrap; (iii) larvae, adults or eggs are found in traps placed at a mutual distance larger than 500 m (Ministère des Solidarités et de la Santé, 2019).

### Entomological longitudinal data

We based our analysis on ovitrap data from three different sources, *i.e.* the VectAbundance, the EID and the Altopictus data repositories. The VectAbundance repository (Da Re et al., 2024) is a standardized dataset of ovitrap records from various locations in Italy, France, Switzerland and Albania covering the period from 2010 to 2022. These records have been homogenized via temporal downscaling on a weekly basis and spatially aggregated on a 9 km grid using the median value of all ovitraps included in such grid box for each observation date. The Altopictus (a French private vector control operator) ovitrap collection includes the municipalities of Rennes (Bretagne region), Bayonne and Saint-Médard-en-Jalles (Nouvelle Aquitaine), Toulouse, Pérols and Murviel-les-Montpellier (Occitania). It is based on a varying number of georeferenced traps (between 15 and 44) that were inspected every 1-2 weeks between 2023 and 2024. The EID (officially “Entente Interdépartementale pour la Démoustication du Littoral Méditerranéen”, a public vector control operator operating along the French Mediterranean littoral) ovitrap collection covers the Provence-Alpes-Côte d’Azur region. It includes the municipality of Nice, based on a varying number of georeferenced traps (between 30 and 50) inspected every 2-4 weeks between 2008 and 2023, and Cagnes-sur-Mer, based on ∼15 georeferenced ovitraps inspected every week between 2011 and 2012 (Lacour et al. 2015).

In order to homogenize ovitrap data available from France with the VectAbundance data, records have been aggregated using the median value for each date.

### Epidemiological data

We retrieved epidemiological data about autochthonous cases of dengue in Western Europe from ECDC (ECDC, 2024c). Data are available from 2010 to 2022 at NUTS3 administrative level.

### Meteorological and environmental data

The dynamic mosquito model is driven by daily meteorological inputs (Metelmann et al., 2019). We retrieved daily rainfall *R* (mm), daily mean *T̅*, maximum and minimum temperatures (°C) from two main sources: E-OBS gridded dataset (at 0.1°) and MétéoFrance weather stations. We approximated the instantaneous temperature *T* based on daily minimum and maximum temperatures using the function employed in Metelmann et al. (2019). E-OBS data were obtained for the period from 2001 to 2023 (Cornes et al., 2018) and its grid was intersected with the human population density raster at 30” based on the GPWv4 data for 2015 (Doxsey-Whitfield et al., 2015) over Western Europe (36°N-60°N, 15°W-19°E) to run simulations. Since VectAbundance and EOBS 0.1° datasets are arranged on a different grids, we used E-OBS at a 0.25° resolution. When cells in the 0.25° grid covered more than one VectAbundance record, we retained only the longest record. For simulating ovitrap data in France, we retrieved weather data from the MétéoFrance weather stations closest to each field site.

We derived the photoperiod (in hours) using the *getSunlightTimes* function from the R package *suncalc*, based on the latitude, longitude and date for each input.

### Model description and experiments

The mechanistic model divides mosquito population into five stages (eggs *E*, diapausing eggs *E*_*d*_, larvae and pupae as juveniles *J*, unfed immature female adults *I*, female adults *A*) and expresses the dynamics of their density (mosquitos/ha) via ordinary differential equations (Figure 1). These equations specify the environmental- and weather-driven transition rates from one life stage to the next:

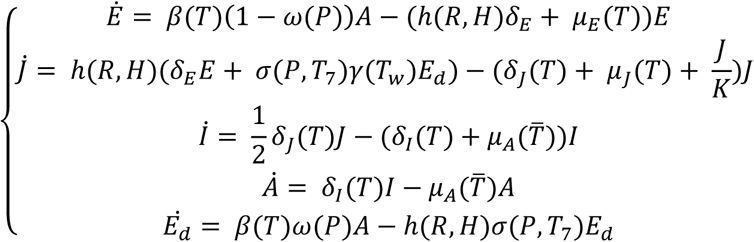

**Figure 1.**
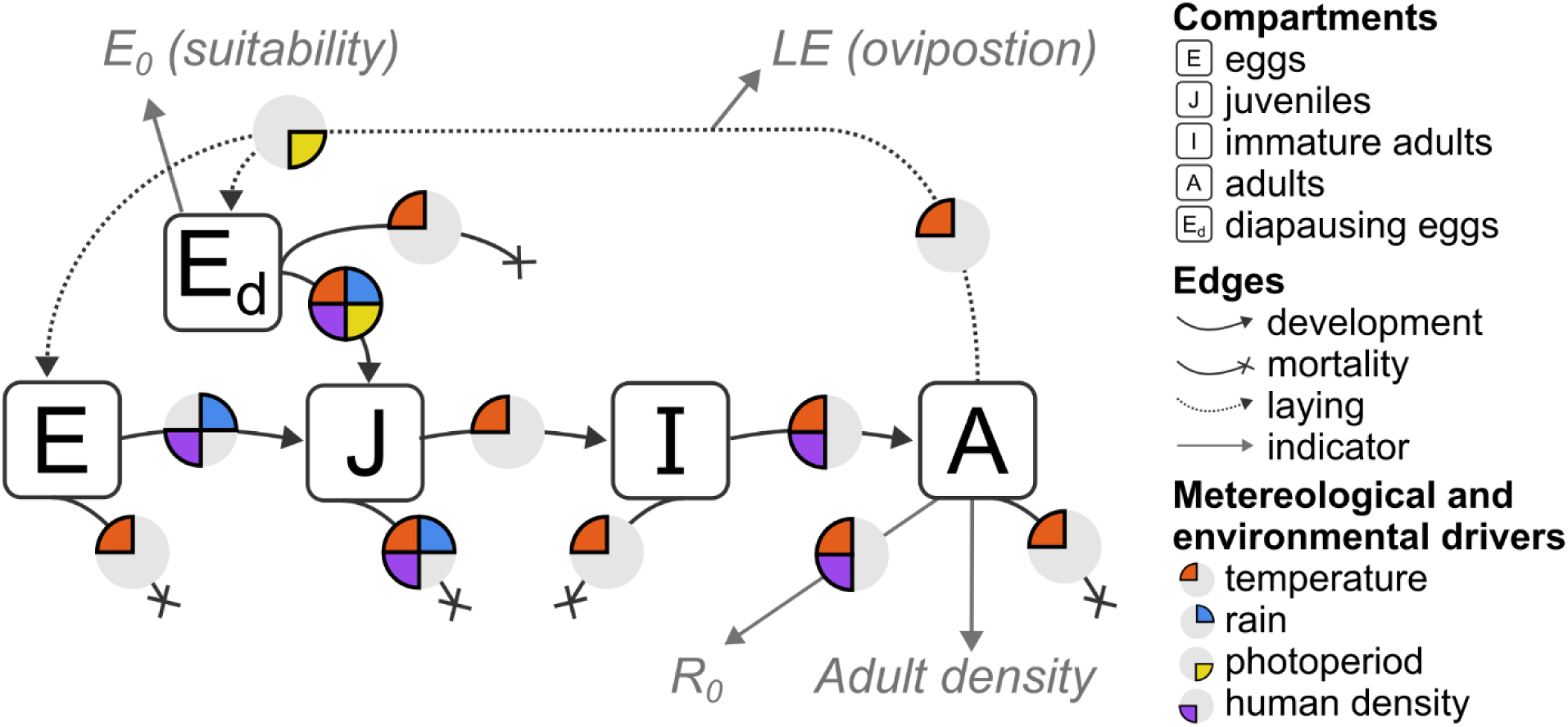
Model outline, as depicted by Metelmann et al. (2019): adults lay eggs which will either hatch or go into diapause depending on the photoperiod. Diapausing eggs will exit diapause because of favorable conditions in terms of temperature and photoperiod. Hatching is due to the presence of water, from rainfall or human irrigation. Juvenile survival depends on the presence of water as well. Remaining development, mortality and egg-laying rates depend on temperature. Immature adults become adults after the first (human) blood meal. Meteorological and environmental drivers are expressed in terms of physical quantities; their attribute is specified in the previous equations.

Development rates of larvae and pupae *δ*_*J*_(*T*) and of immature adults *δ*_*I*_(*T*) (but not of eggs *δ*_*E*_) depend on the instantaneous temperature *T*, as well as the mortality rate of eggs *μ*_*E*_(*T*) and of juveniles *μ*_*J*_(*T*) and the fertility rate *β*(*T*). Adult mortality rate *μ*_*A*_(*T̅*) instead depends on mean daily temperature *T̅*. Production of diapausing eggs is regulated by the parameter *ω*(*P*), which depends on the comparison between the daily photoperiod *P* and a latitude-dependent critical photoperiod (assuming a latitudinal variation as observed in North America; Urbanski et al., 2012), while hatching of diapausing eggs *σ*(*P*, *T*_7_) depends on both photoperiod *P* and mean weekly temperature *T*_7_, and their survival *γ*(*T*_*W*_) on the minimum winter temperature *T*_*W*_. Eggs hatching *h*(*R*, *H*) and the density dependent mortality rate of juveniles, regulated by the carrying capacity *K*(*R*, *H*), depend on the water availability of both natural and anthropic origin, and consequently rely on daily precipitation *R* and fixed human density *H* (details about model assumptions and estimation of thermal traits are available in Metelmann et al., 2019, and Table SI1).

### We run the model under two settings

*Climatic Suitability Setting* – we focus on the temporal changes of spatial climatic suitability, as estimated by the E₀ indicator. The egg population is re-initialized each year on January 1^st^ to 1 ha^-1^ to allow independent assessment of climatic suitability for each year.

*Transient Dynamics Setting* – we estimate temporal changes in adult density, oviposition and R₀ for dengue. The egg population is initialized only once at the start of the simulation (01/01/2006), running a preliminary 10-years spin-up simulation, with no further re-initialization. This setting captures cumulative effect of environmental conditions over time.

### Spatial climatic suitability

We estimated the suitability of a particular location for a given period for *Ae. albopictus* by using the E_0_ indicator, introduced by Metelmann et al. (2019) as an analogous to R_0_ for invasive species. E_0_ is defined as the “expected number of diapausing eggs at the end of a year after having initialized a population with one diapausing egg at the beginning of the same year”. This value is averaged (using the geometric mean) over multiple years for the same site. If, for that site, E_0_ > 1, the growing rate is positive and we assume the site to be suitable for the establishment of *Ae. albopictus* population. Conversely, E_0_ < 1 indicates an unsuitable site.

In order to validate the model, we compared mean E_0_ values in 2018-2023 with observed presence (GBIF.org, 2024) and absence (ECDC, 2024b) data for Europe. Observed data was re-gridded at the same spatial resolution as the EOBS driving climate dataset (0.1°). We then estimated the spatial accuracy of the model using the Area Under the curve (AUC) of the Relative Operating Characteristics (ROC) score (Petrić et al., 2021).

We compared the observed distribution of colonized French municipalities by the “*Direction générale de la santé”* (interpolated at 0.1°) with the model predictions year by year. Simulated establishment of *Ae. albopictus* population was estimated using the aforementioned criterion (E_0_ >1), considering a geometric average over the previous 6 years. We classified the goodness of the classification using the Cohen’s K score (Metelmann et al., 2021). We also calculated the spread rate of *Ae. albopictus* (km/y) in observations and simulations. Spread rate was estimated in each grid cell using the thin plate spline regression of the year of colonization combined with trend surface analysis (Kraemer et al., 2019; Tisseuil et al., 2016).

### Entomological and epidemiological indicators

We simulated the abundance of *Ae. albopictus* at each life stage, the number of laid eggs (per day) and the basic reproduction number R_0_ in Western Europe over the period 2006-23.

The number *LE*_*it*_ of eggs laid per hectare per day in cell *i* on day *t* was estimated using the following:

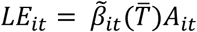

Where *A*_*it*_ is the simulated density of adult female mosquitoes and 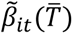 is the approximated fertility rate, defined on mean daily temperature *T̅*. In order to compare our simulations with observed ovitrap data, we averaged *LE*_*it*_ with a moving window of one week (which is the standard trapping period in the VectAbundance dataset), and normalized both simulated and observed laid eggs with respect to maximum value of each trajectory within the time periods of the ovitrap data record. For each time series, we estimated Pearson correlation coefficients and their statistical significance using a Student t-test.

We computed the basic reproduction number, R_0_, for dengue in Europe for each cell *i* and time *t* using the following formulation (Blagrove et al., 2020; Caminade et al., 2017; Metelmann et al., 2021; Zardini et al., 2024):

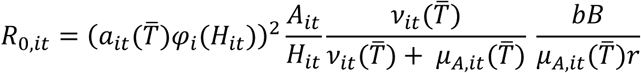

Where (*a_it_* (*T̅*) *φ*) is the biting rate *a_it_* (day^-1^) adjusted for vector preference 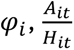 is the vector-to-host ratio, *v*_*it*_ is the reciprocal of the EIP for dengue (day^-1^), *r* is the human recovery rate for dengue (day^-1^), *μ*_*A*,*it*_ (*T̅*) is the aforementioned adult vector mortality rate, *b* and *B* are the probability of transmission of the disease from an infected vector to a susceptible host and *vice versa*. Parameters *a*_*it*_ (*T̅*), *v*_*it*_ (*T̅*) and *μ*_*A*,*it*_ (*T̅*) depend on daily temperature (*T̅*), *φ*_*i*_ depends on human density *H*_*it*_ (per km^2^) and *A*_*it*_ depends on time as it is an output of the dynamical model (Table SI1). We considered the number of days for which R_0_ > 1 (the length of the dengue transmission risk season, LTS) as a proxy for the epidemic risk. We then compared observed autochthonous cases of dengue in 2010-2022 in Europe (ECDC, 2024c), interpolated at 0.1°, with the simulated duration of the epidemic risk. We then estimated the skill of our prediction using the AUC ROC.

### Exposure state analysis

To explore the sensitivity of our simulations to rainfall, temperature and population density, we performed a sensitivity analysis by assessing the variation of the model outputs (i.e. suitability and LTS) by using the exposure state analysis. The exposure space is the n-dimensional space obtained by making incremental change to driving environmental variables to which the system’s outputs are most sensitive and which may change in the future (Culley et al., 2016; Figure SI1). First, we selected 5 sites (using simulations driven by E-OBS 0.25° data) along two gradients of host density and types of climate. We chose Paris city center (11,000 hab/km²) and Paris suburbs (500 hab/km²) as sites with similar climates and different population densities, and Montpellier, (France, Mediterranean climate “Csa”) Bilbao, (Spain, oceanic climate “Cfb”) and Augsburg (Germany, humid continental climate “Dfb”) as sites with similar population density but different climates (*sensu* Köppen classification; Beck et al., 2023). For each location we calculated an exposure space obtained by perturbing recent historical climate conditions (2019-2023) considering average temperatures from May to October *T*_*h*_ (using an additive disturbance, from *T*_1_ = *T*_*h*_ − 2 to *T*_2_ = *T*_*h*_ + 8 °C) and cumulative precipitation from May to October *R*_*h*_ (using a multiplicative disturbance from *R*_1_ = 0 to *R*_2_ = 2*R*_*h*_), for which simulated suitability and LTS werz computed. The perturbation interval for temperatures [-2 °C; +8 °C] was set to consider both recent warming in Western Europe (determined *a posteriori* to be around 1-1.5°C in the last twenty years) and future temperature increase (up to +5/6 °C in the SSP5-8.5 scenario by Carvalho et al., 2021) with an additional buffer.

**Table 1.** Glossary.

| Term | Definition |
| --- | --- |
| Epidemic risk | Epidemic risk is defined as the interaction between a hazard (the presence of an infected virus), the exposure (the contact between an infected vector and a human) and the vulnerability (the probability of developing a disease after a contact). Here we measure it as the length of the transmission risk season (LTS), *i.e.* the number of days for which the basic reproduction number $R_0 > 1$. |
| EIP | The extrinsic incubation period is the time needed for a virus to develop in a mosquito until it reaches the salivary glands, making the mosquito infectious after an infected blood meal. |
| Climatic suitability | Capacity of a site to host a stable mosquito population based on its climate, here defined by $E_0 > 1$ for a given period. $E_0$ has been developed by Metelmann et al. (2019) as an analogue to $R_0$ for invasive mosquito species. It is a measure of the climate-driven yearly net growth rate of a mosquito population in a specific site. |
| Autochthonous (transmission) case | One person locally infected, in opposition to an imported case (infected abroad). For dengue, Zika or chikungunya, a person that is found positive to any of these arboviruses without having visited regions where these arboviruses are endemic. |
| Ovitrap | A trap designed to mimic a development site for collecting eggs laid by a local mosquito population to estimate their oviposition activity. |
| (Immature) development site | A small, natural or man-made, water reservoir suitable for the aquatic life cycle of the mosquito (from eggs hatching to larvae, pupae and the emergence of immature adults). |
| ROC AUC | The Area Under the Curve of the Relative Operating Characteristics is a metric that quantifies the agreement between a presence-absence dataset and a modelled continuous variable (such as $E_0$). |
| Cohen's K | It is a metric that quantifies the accuracy of a prediction of a presence-absence model against a presence-absence dataset which makes full use of the confusion matrix (legend in Figure 3). |
| Thin plate spline regression | An interpolation method of bivariate data. |
| Trend surface analysis | A method to estimate the spatial gradient over an interpolated surface. |
| Exposure space | The n-dimensional space obtained by making incremental change to attributes of environmental variables to which the system's performance is most sensitive and which may change in the future (Culley et al., 2016; details in Figure SI1). |
| NUTS3 | Third administrative level of the Nomenclature of Territorial Units for Statistics of the European Union. |

## Results

### Current climatic suitability for *Ae. albopictus* and risk of autochthonous dengue transmission in Western Europe

Suitable areas for the establishment of the Asian tiger mosquito, as estimated by E_0_, correspond to the lowland and coastal areas in Italy, the Iberian Peninsula, central and southern France, and the Adriatic Balkans (Figure 2a). Climate is also suitable over northern European cities such as Paris, London, Vienna, Strasburg and Frankfurt. Observed occurrence of *Ae. albopictus* (GBIF.org, 2024) roughly follows a similar pattern, with several false positives (i.e., areas simulated to be colonized, but uncolonized according to the observations) over the inner southern Iberian Peninsula. Despite this regional discrepancy, the suitability for establishment reproduces well the current distribution of *Ae. albopictus* in Western Europe (ROC AUC = 0.90).

**Figure 2.**
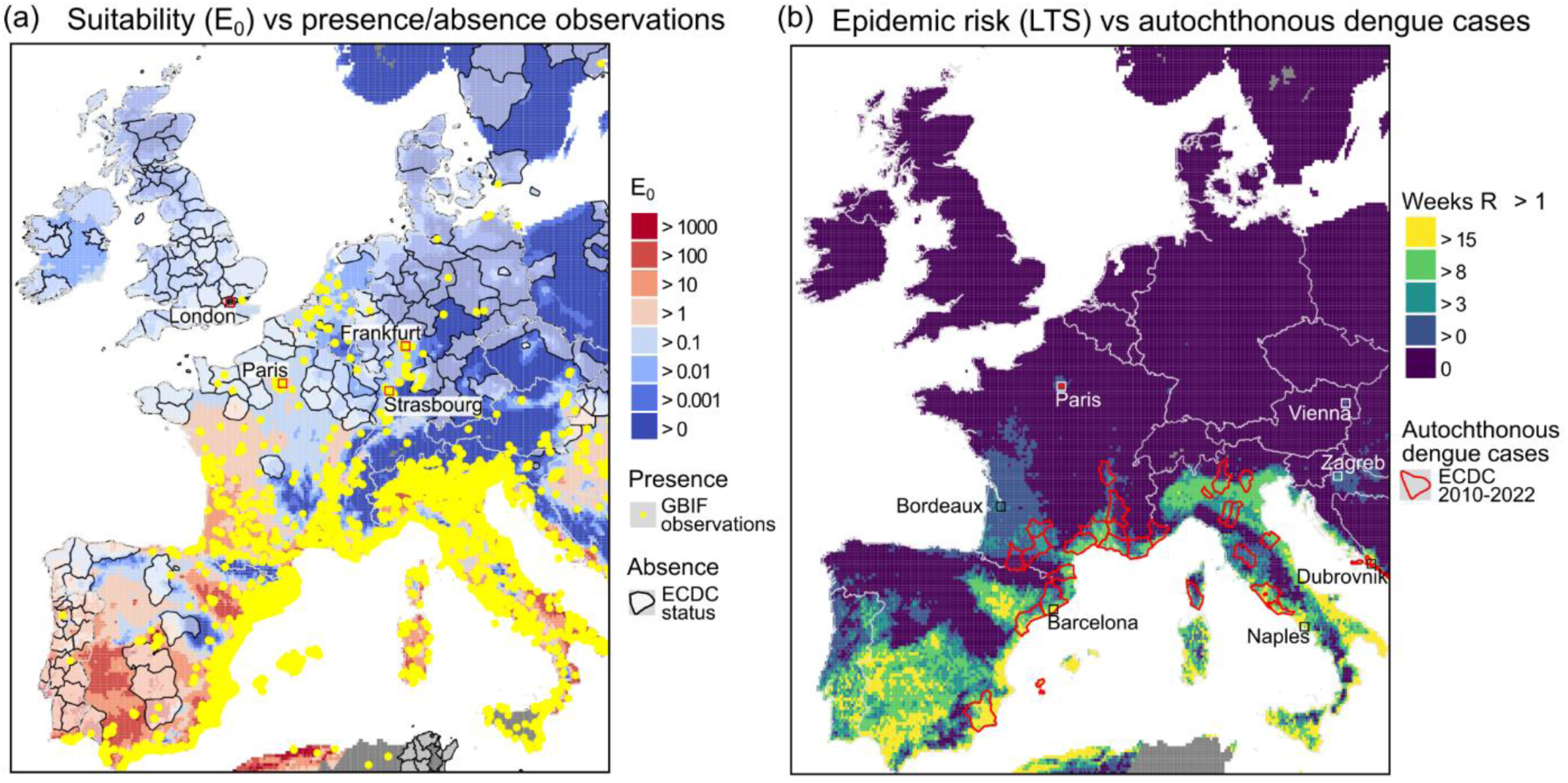
Suitability of Western Europe for the establishment of Ae. albopictus and epidemic risk of dengue compared to observations. (a) Geometric average of E_0_ in 2018-2023, GBIF observed occurrence data (yellow dots) and absence data (white transparent layer) from ECDC. (b) LTS, i.e. the annual mean duration of the dengue transmission season in 2010-2022, (defined as the period for which R_0_ > 1), and autochthonous dengue cases as reported by ECDC at the NUTS3 level. Missing values due to unavailability of weather data are depicted by the gray shading.

Areas potentially at risk of dengue transmission correspond to a subset of areas suitable for *Ae. albopictus* (Figure 2b). The highest LTS values are simulated over coastal areas along the Mediterranean and Adriatic Sea, where R_0_ > 1 for more than 3 months on average (e.g., LTS in Barcelona is 18.1 weeks, Naples 15.8 weeks, Dubrovnik 14.8 weeks). Over the Atlantic coasts of France and Spain, this period is shorter and ranges between 1 and 3 weeks per year (e.g. Bordeaux 2.9 weeks). Autochthonous transmission of dengue might also be possible in large urban cities such as Zagreb (LTS = 3.0 weeks), Vienna (LTS = 1.1 weeks) and Paris (LTS = 1.0 week).

Observed autochthonous cases of dengue were mostly reported in southern France, Italy, Croatia and over the eastern coasts of Spain. The performance of LTS to predict autochthonous cases is moderate (ROC AUC = 0.75).

### Temporal analysis of the colonization of France by *Ae. albopictus*

The colonization of France by the Asian tiger mosquito started from the southeastern regions, namely in Alpes-Maritimes in 2004 (Roche et al., 2015). The model predicts a wider area to be already suitable for *Ae. albopictus* in 2006, around the Mediterranean coasts, the southern Atlantic coasts and cities like Paris and Lyon (Figure 3a). This spatial mismatch between newly colonized areas and the simulated environmental niche leads to a poor model performance (Cohen’s K = 0.02; Figure 3e). During the following years, both suitable and colonized areas have gradually expanded (Figure 3b-c-d). The observed colonization front accelerated roughly from 10 to 20 km/y (median value) in 10 years, while climatically suitable areas expanded at a slower rate, less than 5 km/y, with an acceleration simulated over the last 5 years (Figure 3f). Therefore, the model skill increases with time (Cohen’s K > 0.5 since 2014, with a maximum of 0.64 reached in 2018). A larger geographical domain was already suitable for the establishment of *Ae. albopictus* when it first colonized Nice and parts of Corsica in 2006 and that *Ae. albopictus* has gradually invaded its theoretical climatic niche over the 2006-2024 period. We observe an overestimation of suitable areas over the western regions and an underestimation over mountain fringe areas in Corse, the Alps, the Pyrenees and the Massif Central, but also in Alsace (Figure 3d).

**Figure 3.**
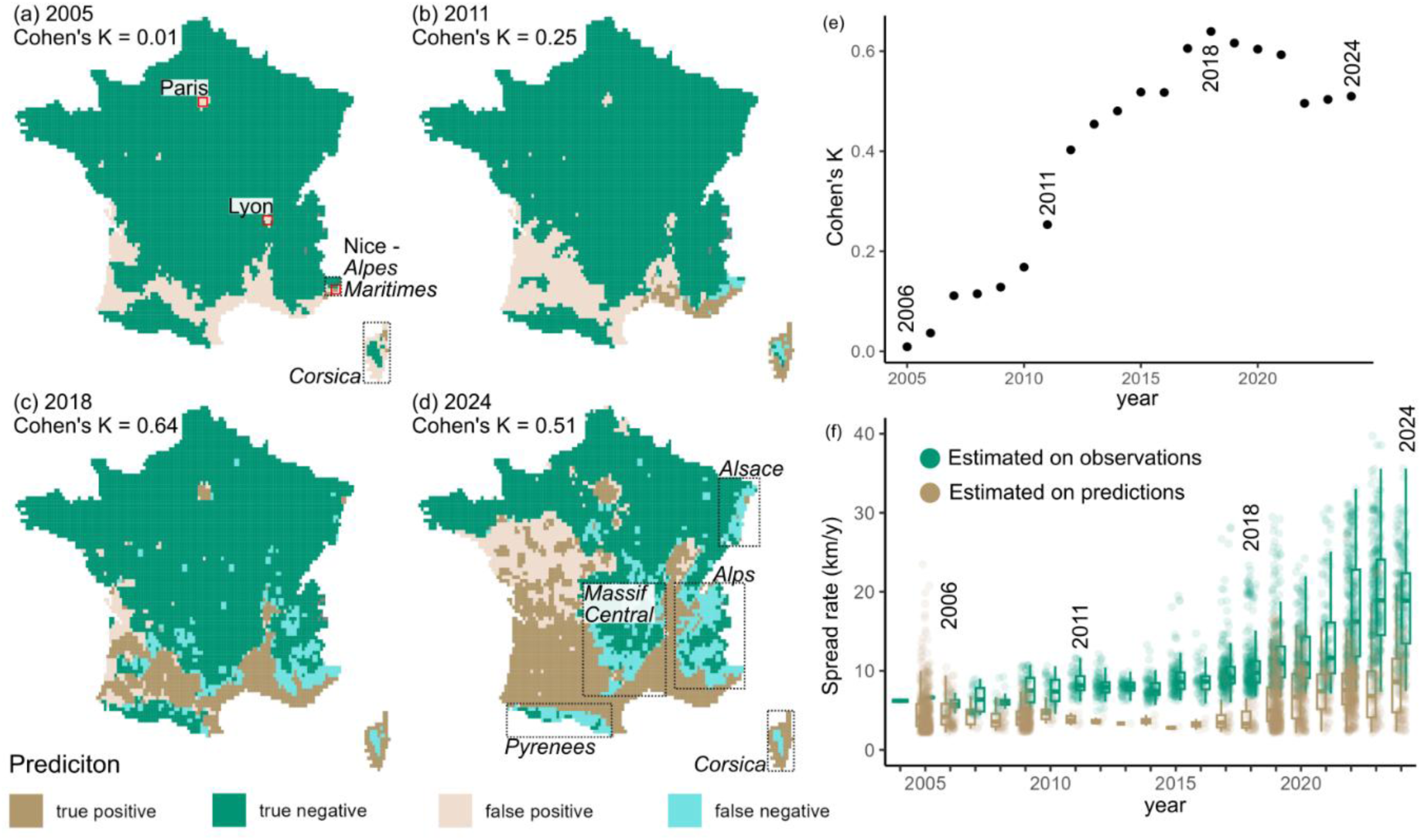
Comparison between the observed colonization of Ae. albopictus in metropolitan France (a-d) and simulated suitability niche (E_0_ > 1) between 2005 and 2024, (e) model accuracy (Cohen’s K) and (f) spread rate (km/year) estimated from both observations and predictions.

### Estimation of oviposition activity

We simulated oviposition activity for 74 sites in France, Switzerland and the Italian peninsula. We observed a positive linear correlations between simulations and ovitrap observations for 70 sites (see Figure SI2). 66 out of 74 correlation coefficient values were statistically significant at the 99% confidence interval (p-value < 0.01 based on a Student T-test, Figure 4a). The seasonal cycle of simulated egg abundance was well captured by the model for some locations (Figure 4b-e). The lowest correlations are shown over the northeastern part of Italy.

**Figure 4.**
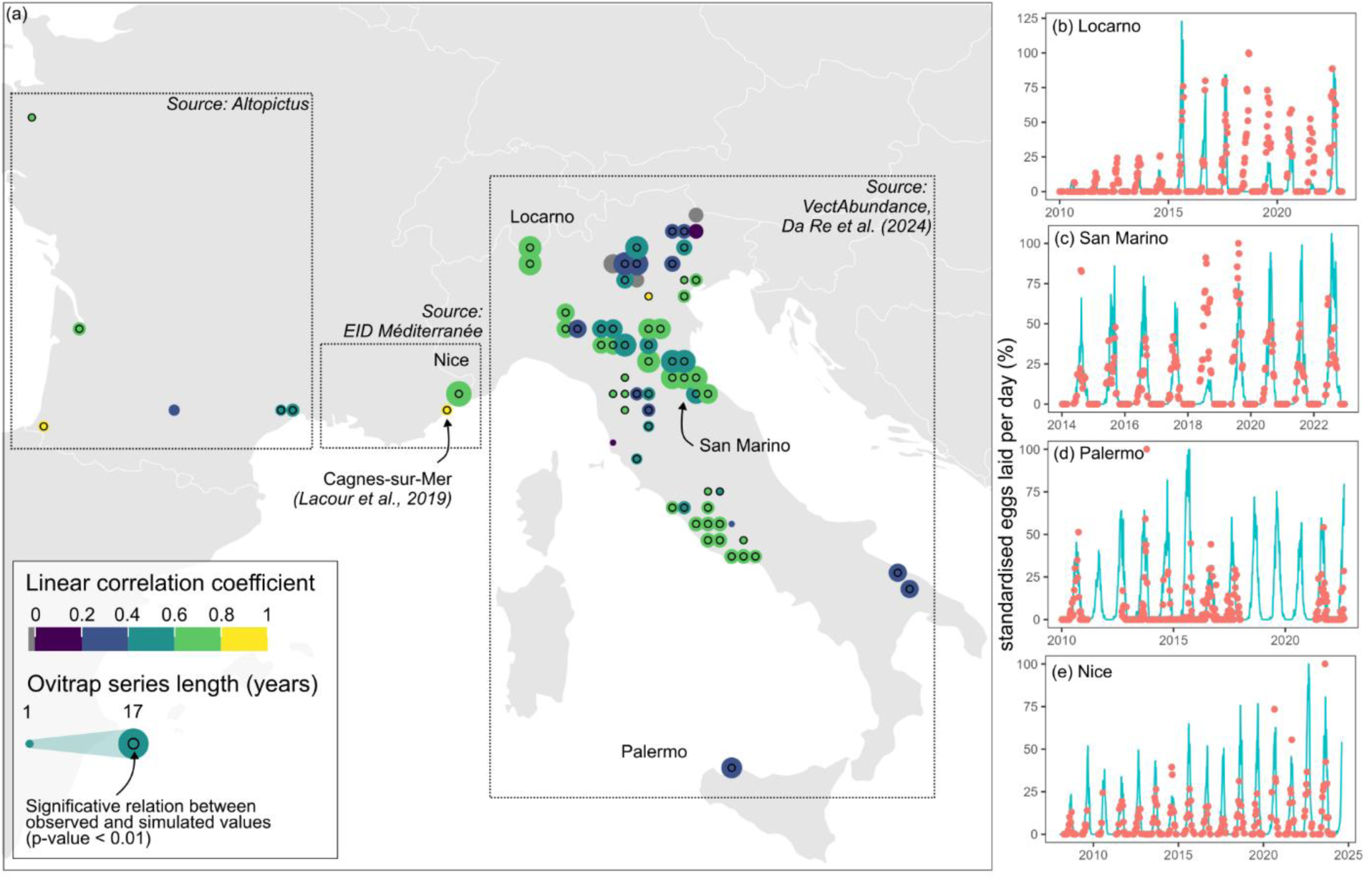
Comparison between observed (orange dots) and simulated (blue line) standardized laid eggs rate per day (%). (a) Localization of ovitraps and Pearson linear correlation coefficient values with model predictions. Comparison between simulations and observations for four different locations: (b) Locarno (CH, r: 0.58), (c) San Marino (SM, r: 0.58), (d) Palermo (IT, r: 0.37), (e) Nice (FR, r: 0.66). The other time series are reported in Figure SI2. Ovitrap data was mainly derived from VectAbundance (Da Re et al., 2024). Since the normalization is done with respect to the maximum value within the record period, simulation values over 100% are admissible outside the record period (such as in Locarno).

### Effect of recent climate change conditions on simulated suitability, adult density and dengue epidemic risk

For estimating the impact of recent climate change conditions, we compared two periods: 2006-2014 (“2010s”) and 2015-2023 (“2020s”). By the 2010s, a large part of southern Europe was already suitable (E_0_ > 1) for the establishment of the Asian tiger mosquito, with the exception of mountainous areas (such as the Pyrenees between Spain and France or the Apennines in Italy; Figure 5a). During the 2020s, an increase in E_0_ is simulated over Western Europe (Figure 5b). Importantly, 2020s climatic conditions are simulated to become suitable for the establishment of *Ae. albopictus* over the Western lowlands of France, over the central plateau of Spain, and in large urban cities such as London, Vienna and Frankfurt. The suburbs of Paris and Zagreb are also simulated to become increasingly suitable during the 2020s (Figure 5c).

**Figure 5.**
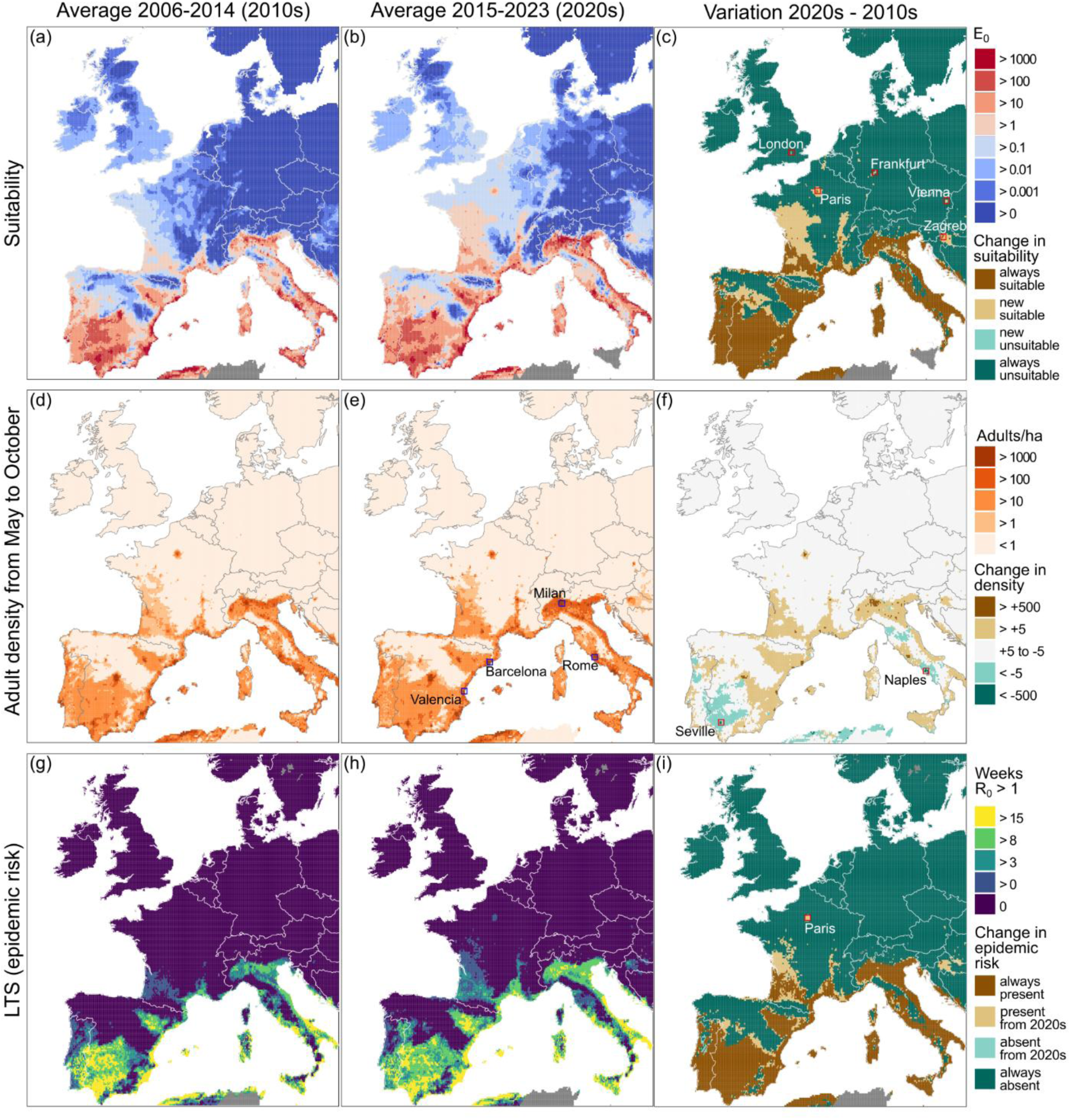
Simulated effect of recent climatic conditions on the suitability of Ae. albopictus (upper row), adult density during the activity period (middle row), and on dengue transmission risk (bottom row) over Western Europe. (a) Suitable areas, measured using E_0_ in 2006-2014, (b) in 2015-2023 and (c) variation between the two periods (considering the threshold E_0_ >1). (d) Average adult abundance (per ha) between May and October for the period 2006-14, (b) 2015-23, (c) and variation. (g) LTS for dengue (number of days in which R_0_ > 1) for 2006-14, (h) 2015-23 (i) and variation.

Changes in simulated adult density (per ha) follow a similar spatial pattern. There is an increasing trend in simulated adult density over the southern part of Western Europe and Paris during the 2020s (Figure 5e). Interestingly, a slight decrease in adult density is simulated over large areas over the southernmost part of Europe, particularly over southern Spain (Sevilla) and southern Italy (Naples; Figure 5f). Highest simulated densities of mosquitoes are reached in Barcelona (23,000 adults/ha), Valencia (19,000 adults/ha), Milan and Rome (15,000 adults/ha).

Most of Western Europe is still unsuitable for autochthonous transmission of dengue for the 2020s (Figure 5h). However, the LTS is higher than 2 months on average over most of Italy, south-eastern France, Spain and Portugal. Largest values (> 3 months) are simulated in coastal cities bordering the Mediterranean and Adriatic Sea. Simulated LTS for dengue show large values in Italy, Spain and Portugal (maximum values up to 37 weeks in Sicily and Sardinia, 29 in southern Portugal) and coastal areas in Croatia (up to 18 weeks) during the 2010s (Figure 5g). New regions at risk (LTS >0) emerge in 2020s over most of Central-Western Europe, while LTS decreases over the warmest regions of the southern Iberian Peninsula and Italy (consistently with the simulated decrease in adult density). Importantly, Nouvelle Aquitaine region in France, parts of central Spain, inner Croatia and Paris become suitable for autochthonous transmission of dengue during the 2020s (Figure 5i).

### Model sensitivity to rainfall, temperature and human population density

By using the exposure space analysis, we assessed the influence of temperature, precipitation variations and host population density gradients on *Ae. albopictus* dynamics and dengue transmission risk in Western Europe. Regarding host density, we considered the two Parisian sites along a decreasing host population gradient with an oceanic climate “Cfb” of the Köppen classification (Beck et al., 2023). In the exposure space, we traced the 1.3 °C increase in average temperature from May to October, with no clear changes in annual rainfall between 2004 and 2023 in Paris (gray lines on Figure 6a-d). Simulated E_0_ is larger in the densely populated area of Paris (direct impact of carrying capacity that depends on rainfall and human host density in the model). Paris center has been suitable for the whole study period (Figure 6a), while suburbs became suitable only during recent years with recent increases in temperature (Figure 6c). Concerning the dependency of E_0_ to rainfall, contour lines tend to be rounded – rather than straight vertical – while decreasing in population density (Figure 6b, d). In densely populated areas, rain plays a minor role, due to permanent man-made development sites, whereas in sparsely populated areas rain becomes a limiting factor. The effect of host density on LTS is more complex, since a higher host abundance also reduces the vector-to-host ratio in simulated R_0_. For such a reason, the suburbs of Paris are simulated to experience a higher risk of dengue transmission at higher temperatures (Figure 6d) compared to the city center (Figure 6b).

**Figure 6.**
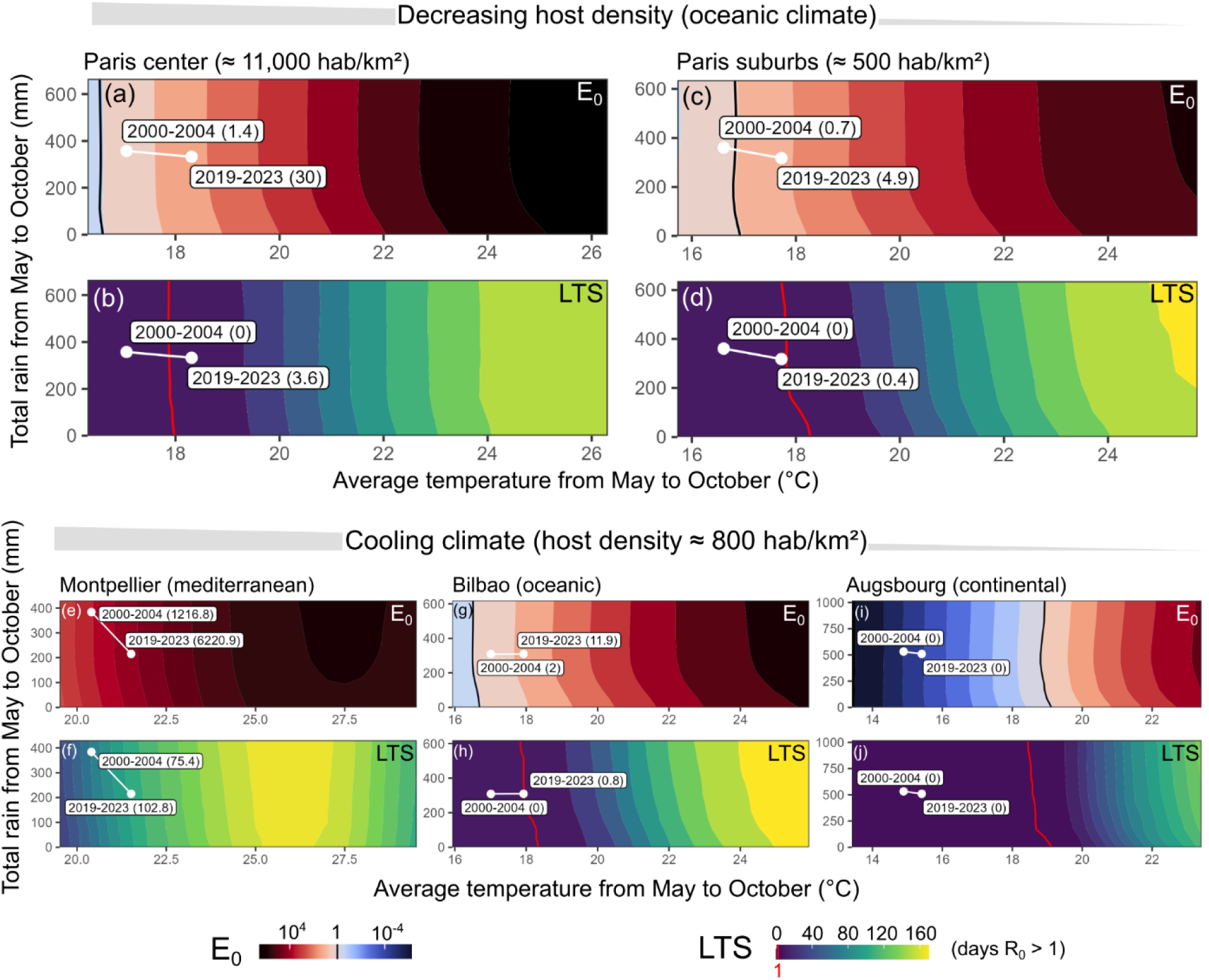
Exposure spaces generated by perturbing historical (2019-2023) weather trajectories of temperature (°C) and precipitation (mm) in six cities of Western Europe. For readability, the x-axis shows the average temperature from May to October, while the y-axis the cumulative precipitation. For each location, we plot the climatic suitability as measured by E_0_ (top panel with a black line denoting threshold E_0_ = 1) and the length of the dengue transmission season (in days, bottom panel with a red line depicting the threshold R_0_ = 1). In Montpellier, E_0_ and LTS are greater than 1 in the whole exposure space. In each figure, we show the climatic trajectory of each site over the last 20 years.

We also compared three different types of climate on simulated E_0_ and LTS while keeping similar population density values (∼800 hab/km²). The Mediterranean climate in Montpellier is highly conducive for the establishment of *Ae. albopictus* population. However, a gradual decrease in all simulated indicators is shown when temperatures exceed about 27°C (Figure 6e, f). For Bilbao, a southern European oceanic city, climatic conditions became suitable for disease transmission in 2023 (red line in Figure 6h). Recent humid continental climate conditions, as experienced in Augsburg, southern Germany, are still unsuitable for the vector and potential risk of dengue transmission (Figure 6i,j).

## Discussion

### Climate change is key to understand the recent spread of the Asian tiger mosquito in Western Europe

The ongoing invasion of *Ae. albopictus* is increasingly raising concerns about the future of tropical diseases in Europe. 2024 represents a record year in terms of dengue cases at global scale (about 14,000,000, mostly reported in Latin America; ECDC, 2024c), as well as autochthonous cases of arboviruses in metropolitan France (84; Santé Publique France, 2024) and the largest dengue outbreak ever reported in Western Europe (199 in Fano, Italy; Sacco et al., 2024).

In this context, our study aimed to (i) validate simulations with respect to an ensemble of entomological and epidemiological observations and (ii) test the hypothesis that recent climate change affected the establishment of *Ae. albopictus* population and its potential to transmit dengue in Western Europe. We confirmed the skill of the E_0_ indicator (AUC = 0.90) in matching the distribution of *Ae. albopictus* in Western Europe. The main discrepancy between observations and model predictions occurs over the southern Iberian Peninsula, where high suitability is simulated (as in other studies, such as Zardini et al., 2024), though *Ae. albopictus* has not been reported yet. Such mismatch could be considered further by using rainfall thresholds to calculate aridity conditions in our modeling framework. Former studies based on threshold-based models show that at least 400-500 mm of total annual rainfall is needed for the survival of the Asian tiger mosquito (Kobayashi et al., 2002; Petrić et al., 2021). However, *Ae. albopictus* can lay its eggs in man-made water reservoirs, which will play an important role regardless of rainfall amounts (Paupy et al., 2009; Tran et al., 2013). Alternatively, the absence of mosquitoes in the region may be explained by the slow process of arrival, adaptation and colonization of the region (Barman et al., 2024). As a matter of fact, *Ae. albopictus* was first detected in Portugal in 2017 (Osório et al., 2018), surprisingly late compared to other southern European countries. Such delay might be related to passive transportation factors and geographical barriers that might have prevented an earlier introduction of this species in Portugal (Zé-Zé et al., 2024).

Understanding colonization dynamics is key to interpret the observed establishment of *Ae. albopictus* in France, as well as estimating its spread in the upcoming years. The colonization of a new region by an invasive species results from the interplay between *i*) the mobility of the invasive species across a suitable area, *ii*) its adaptation to new environments and *iii*) the expansion of suitable areas. Importantly, the skill-score we used to estimate the agreement between our climatic suitability model and observed colonization of French municipalities was high in 2018 (Cohen’s K = 0.64) but close to zero in 2006 (Cohen’s K = 0.02). We propose that the systematic overestimation of model predictions before 2010 unravels the gradual colonization of an already climatically suitable area, very likely driven by passive motor vehicle transportation of the vector (Roche et al., 2015). Therefore, the improved model performance in recent years (Cohen’s K ∼ 0.5, Figure 3e) reflects a progressive overlap between the simulated and the observed niches, whose conceptual distinction is crucial in forecasting climate effects on ectotherms (Braschler et al., 2020). This suggests that *Ae. albopictus* is approaching the limits of its theoretical climatic niche, and that future colonization may be driven by the climate-induced enlargement of the suitable area. However, one cannot overlook instances of colonization outside these areas (e.g. in Alsace, Figure 3d), where either local populations may be adapting to colder climates or environmental drivers of suitability are poorly captured by the model, and might require further investigation. The accelerating colonization of *Ae. albopictus* in France, with a spread rate from 10 to 40 km/year, is qualitatively consistent with findings by Kraemer et al. (2019), who obtained larger values (100 to 200 km/year) based on mosquito occurrence data (Kraemer et al., 2015) for Europe.

### Recent climate change has different impacts on *Ae. albopictus* suitability, its abundance and its potential to transmit autochthonous cases of dengue

Our simulations highlight the expansion of areas suitable for the establishment of *Ae. albopictus* in Europe, consistently with predictions by other studies (Colón-González et al., 2021). These areas, located between 40° and 52°N, are either located over large continuous regions (such as the Spanish plateau and the French Western Atlantic region) or in isolated urban centers (London, Frankfurt, Vienna, and Zagreb). In cities, the urban heat island effect combined with high population density could provide suitable conditions for *Ae. albopictus*, compensating for inadequate conditions of the surroundings. The model predicts a higher density of adults compared to other estimates (Brass et al., 2024 and Zardini et al., 2024) with a more heterogeneous distribution, e.g., hotspots simulated in cities (compared to estimates by Brass et al., 2024). The predicted geographical expansion of adult presence for the 2020s are similar to estimates by Zardini et al. (2024) and more conservative to other studies (Brass et al., 2024; Petrić et al., 2021) which simulate adult presence in central and northern Europe. Simulated adult density seems to increase in 2020 rather than spreading to new areas, notably between 40° and 49°N. Conversely, in a warm Mediterranean region (between 37° and 43° N), corresponding to the southern Iberian Peninsula, southern and central Italy, we estimate a localized decrease in simulated adult density. Such decrease may be related to summer temperatures exceeding optimal values for the survival of adult mosquitoes (around 26-27 °C in the model; Metelmann et al., 2019) as well as decreasing precipitation with a detrimental effect on oviposition. A recent modeling study confirms that heat wave have contrasting effects on mosquito abundance in temperate climates in Italy – positive in northern continental areas and detrimental in southern warmer ones (Garrido Zornoza et al., 2024). Estimating the effect of climate change on the risk of dengue transmission is more challenging, since one needs to consider the viral development inside the ectotherm vector as well as other important human behavior factors (Morin et al., 2013). At low temperatures, the extrinsic incubation period (EIP) is usually too long for the vector to become infectious during his standard life span (about 30 days; Morin et al., 2013). At higher temperatures, there is a competition between shorter EIPs (favoring disease transmission) and mosquito survival (Blagrove et al., 2020). The vector-to-host ratio is also an important factor in R_0_ formulation. It can lead to counterintuitive outcomes, since higher host density typically leads to increased carrying capacity (with a saturating effect for very high human density – a factor neglected by the model) and increased vector abundance, but also lower vector-to-host ratio values. We found that the sensitivity of epidemic risk to temperature increase is highest in middle-sized cities. This theoretical result is consistent with recent observations. An outbreak of chikungunya was reported in Ravenna (about 150k hab.) in 2007 (Angelini et al., 2007) and an outbreak of dengue affected Fano (about 60k hab.) in 2024 (Sacco et al., 2024). Simulations predict suitability for the transmission of dengue along the Mediterranean coasts and new areas (mainly in France) to become suitable in the 2020s. This finding is globally consistent with the distribution of reported cases of dengue in Croatia, eastern Spain, southern France, Paris, and the Italian peninsula (ECDC, 2024c). Overall, simulated LTS increased by 3.6 days between the 2010s and the 2020s for the study area; this is qualitatively consistent with the projections by Colón-González et al. (2021), who estimated that Europe (>1500 m.a.s.l.) might experience a 1-3 weeks increase in LTS in 2006-2099 compared to 1951-2006, depending on altitude and host density.

### Caveats and future directions

The model we employed provides a framework for describing environmentally driven mosquito population dynamics. It is a parsimonious framework that could be refined by incorporating genetic and phenotypic plasticity (Brass et al., 2024), delayed biological effects (Metelmann et al., 2021), the distinction between parous and nulliparous females (Tran et al., 2013), human mobility (Barman et al., 2024; Da Re et al., 2022), the relationship between the human-influenced landscape and the carrying capacity of the juvenile stage (Lamy et al., 2023). However, this model takes into account the latitudinal variation of the critical photoperiod that controls the hatching of overwintering diapausing eggs. This adaptation feature, which triggers egg hatching in spring when temperatures are more favorable, was observed in North America already twenty years following the introduction of *Ae. albopictus* (Urbanski et al., 2012). We argue that evolutionary convergence might have caused the same adaptation behavior in *Ae. albopictus* populations over the European continent (Lacour et al., 2015). Yet, other types of adaptation could occur, such as an increased resistance to warm temperatures once they become the limiting factor (Couper et al., 2021), or that winter diapause might even become unnecessary for the persistence of *Ae. albopictus* populations in warm temperate regions. Without further experimental studies about adaptation mechanisms, future predictions of *Ae. albopictus* mostly rely on historical traits and consequently remains uncertain. The future of dengue in Western Europe depends also on the adaptation of another competent vector of arboviruses, *Ae. aegypti*, which is known to be less tolerant to temperate climates compared to *Ae. albopictus* (Souza & Weaver, 2024). *Aedes aegypti* is currently absent in Western Europe (ECDC, 2024a), but its recent establishment in Cyprus and Madeira (Portugal) suggests that projections of future epidemic risk should also consider this vector. To estimate the length of the dengue transmission season, we used a typical R_0_ formulation (Blagrove et al., 2020; Caminade et al., 2017; Zardini et al., 2024) that is valid at the equilibrium for fixed parameters, for completely susceptible population. Since dengue is not endemic to Europe, we can assume the latter condition to be satisfied. However, R_0_ computation in the case of time-varying parameters may lead to an overestimation in disease risk (Djidjou-Demasse et al., 2020). To overcome this limitation, future modeling work could be based on a SEIR (Susceptible-Exposed-Infectious-Recovered) model to simulate the number of autochthonous cases.

A major caveat in this study is the mismatch in the spatiotemporal resolution among different data sources. In the ECDC dataset of dengue transmission cases, the entire NUTS3 region is marked as ‘positive’, although the transmission event occurred locally (Figure 3d) and our estimation of the epidemic risk is limited by the resolution of the driving weather data (0.1 °). The same applies to the dataset of the colonization in France at the municipality scale, thus resulting in a systematic overestimation of the colonized areas over mountainous areas (in Corsica, the Alps, the Pyrenees, the Massif Central; Figure 2). Furthermore, entomological data may be biased by delays in reporting by field operators. Conversely, the E-OBS weather dataset was too coarse when comparing simulated egg density and observed ovitrap time series in high environmental gradients of alpine valleys (Figure 4). In order to simulate the future dynamics of *Ae. albopictus* using climate model outputs for different SSP-RCP scenarios, following studies will need to consider these discrepancies.

### Recommendations for vector control operators and public health officers

#### Reinforce collaboration between surveillance operators and academic researchers

This work would not have been possible without collecting reliable and up-to-date observation data on mosquito presence and abundance. This highlights the need to maintain strong and continuous links between surveillance operators and academic researchers in terms of optimizing protocols and data sharing. Maintaining and expanding these efforts would ensure the interception of adaptation mechanisms that would lead to drastic system changes (e.g. local vector adaptation) and the improvement of models.

#### Focus entomological surveillance on newly suitable areas for vector establishment

The gap between the simulated and the observed niches of *Ae. albopictus* may be progressively closing. Future entomological surveillance should focus on areas projected to become suitable, independently of the geographic distance with presently colonized areas. Such areas include, but are not limited to, the western French countryside and large, populated cities between Southern England and Western Germany, such as London and Frankfurt.

#### Focus sanitary surveillance on newly suitable areas for arboviral transmission

Countries would benefit from prioritizing surveillance and control measures in areas predicted to be suitable for *Ae. albopictus* or arboviral transmission. This would help refocus anticipatory measures and avoid dispersing resources over low-risk areas – as in France, for example, where the Decree of the 23^rd^ of July 2019 defines the entire national territory as at risk for vector-borne transmission. Moreover, the sensitivity analysis and observations of autochthonous arbovirus cases (Angelini et al., 2007; ECDC, 2024c; Roiz et al., 2015; Sacco et al., 2024; Santé Publique France, 2024) suggest to target prevention and control efforts against arbovirus transmission in mediumly populated areas that were colonized by *Ae. albopictus*.

## Acknowledgements

The authors acknowledge the support of RIVOC funded by the Région Occitanie (France) for the VectoClim project. In addition, A.R. acknowledges precious scientific discussions with Soeren Mettelmann, Ramses Djidjou Demassé, Benjamin Roche, Mircea Sofonea for modelling suggestions, as well as Antoine Mignotte, Frédéric Jean and Daniele Da Re for entomological insights. A.R. acknowledges also the use of the i-Trop HPC and would like to thank the team of the i-Trop platform for their technical support.

