## Supplementary Information for "*Aedes albopictus* is rapidly invading its climatic niche in France: wider implications for biting nuisance and arbovirus control in Western Europe"

### Additional materials

#### Model parameters

In the modelling framework established by Metelmann et al., (2019) both the carrying capability $K_{it}$ and the hatching $h_{it}$ of a site $i$ at time $t$ depend on host density $H_{it}$ and rainfall $r_{it}$ :

$$K_{it} = \lambda \frac{1- \alpha_{evap}}{1- \alpha_{evap}{}^{t}} \sum_{x=1}^{t} \alpha_{evap}{}^{(t-x)} \left( \alpha_{rain} R_{ix} + \alpha_{dens} H_{ix} \right)$$

$$h_{it} = (1- \varepsilon_{rat}) \frac{(1+\varepsilon_0) e^{-\varepsilon_{var}\left(R_{it}-\varepsilon_{opt}\right)^2}}{e^{-\varepsilon_{var}\left(r_{it}-\varepsilon_{opt}\right)^2} + \varepsilon_0} + \varepsilon_{rat} \frac{\varepsilon_{dens}}{\varepsilon_{dens} + e^{-\varepsilon_{fac} H_{it}}}$$

The values of the model's parameters (carrying capacity, vector model and Ro model) and associated references are shown in Table SI1.

*Table SI1 - Model parameters*

| Parameter | Meaning | Value/formula | Source |
|---|---|---|---|
| $CTT_S$ | critical temperature over one week in spring (°C ) | $11.0$ | (Metelmann et al., 2019) |
| $CPP_S$ | critical photoperiod in spring (hours) | $11.25$ | (Metelmann et al., 2019) |
| $\sigma(T,P)$ | spring hatching rate (day$^{-1}$) | $\begin{cases} 0 \text{ if } \mathrm{T_7} < \mathrm{CTT_S} \text{ or } \mathrm{P} < \mathrm{CPP_S} \\ 0.1 \text{ otherwise} \end{cases}$ | (Metelmann et al., 2019) |
| $CPP_A(L)$ | critical photoperiod in autumn (hours) | $10.058 + 0.08965\mathrm{L}$ | (Metelmann et al., 2019) |
| $\omega(P)$ | fraction of eggs going into diapause | $\begin{cases} 0 \text{ if } \mathrm{P} < \mathrm{CPP_A} \text{ or day} < 183 \\ 0.5 \text{ otherwise} \end{cases}$ | (Metelmann et al., 2019) |
| $\delta_E$ | normal egg development rate (day$^{-1}$) | $1/7.1$ | (Metelmann et al., 2019) |
| $\delta_J(T)$ | Juvenile development rate (day$^{-1}$) | $1/(83.85 - 4.89\,\mathrm{T} + 0.08\,\mathrm{T}^2)$ | (Metelmann et al., 2019) |
| $\delta_I(T)$ | first pre-blood meal rate (day$^{-1}$) | $1/(50.1 - 3.574\,\mathrm{T} + 0.069\,\mathrm{T}^2)$ | (Metelmann et al., 2019) |
| $\mu_E$ | egg mortality rate (day$^{-1}$) | $-\ln\left(0.955\,\mathrm{e}^{-0.5(\mathrm{T}-18.8/21.53)^6}\right)$ | (Metelmann et al., 2019) |
| $\mu_J$ | juvenile mortality rate (day$^{-1}$) | $-\ln\left(0.977\,\mathrm{e}^{-0.5(\mathrm{T}-21.8/16.6)^6}\right)$ | (Metelmann et al., 2019) |
| $\mu_A(\overline{T})$ | adult mortality rate (day$^{-1}$) | $-\ln\left(0.677\,\mathrm{e}^{-0.5(\overline{\mathrm{T}}-20.29/13.2)^6} 0.069\,\overline{\mathrm{T}}^{0.1}\right)$ | (Metelmann et al., 2019) |

| | | | |
|---|---|---|---|
| $\gamma(T_w)$ | survival probability of diapausing eggs (winter$^{-1}$) | $0.93\ e^{-0.5(T_w-11.68/15.67)^6}$ | (Metelmann et al., 2019) |
| $\beta(T)$ | egg laying rate (day$^{-1}$) | $\begin{cases} 33.2\ e^{-0.5(T-70.3/14.1)^2}(38.8-T)^{1.5} \text{ if } T \leq 38.8 \\ 0 \text{ otherwise} \end{cases}$ | (Metelmann et al., 2019) |
| $\lambda$ | capacity parameter (larvae days ha$^{-1}$) | $10^6$ | (Metelmann et al., 2019) |
| $\alpha_{evap}$ | Normalization parameter evaporation | 0.9 | (Metelmann et al., 2019) |
| $\alpha_{dens}$ | Normalization parameter of host density (km$^2$) | $10^{-5}$ | (Metelmann et al., 2019) |
| $\alpha_{rain}$ | Normalization parameter of rain (mm$^{-2}$) | $10^{-2}$ | (Metelmann et al., 2019) |
| $\varepsilon_{rat}$ | Proportion of hatching due to rain | 0.2 | (Metelmann et al., 2019) |
| $\varepsilon_0$ | Correcting additive parameter for rain-related hatching | 1.5 | (Metelmann et al., 2019) |
| $\varepsilon_{var}$ | Stretching parameter for rain-related hatching (mm$^{-2}$) | 0.05 | (Metelmann et al., 2019) |
| $\varepsilon_{opt}$ | Correcting additive parameter for human-induced hatching (mm) | 8 | (Metelmann et al., 2019) |
| $\varepsilon_{dens}$ | for rain-related hatching | $10^{-2}$ | (Metelmann et al., 2019) |
| $\varepsilon_{fac}$ | Stretching parameter for human-induced hatching (km$^2$ km$^{-2}$ | $10^{-2}$ | (Metelmann et al., 2019) |
| $a$ | Mosquito biting rate | $(0.0043T + 0.0943)/2$ | (Blagrove et al., 2020; Caminade et al., 2017; |

| | | | |
|---|---|---|---|
| | | | Zardini et al., 2024) |
| $\varphi$ | Human biting preference | $\begin{cases} 0.9 \text{ if human density} > 50 \text{ hab km}^{-2} \\ 0.5 \text{ otherwise} \end{cases}$ | Value of 0.5 by (Caminade et al., 2017) |
| $EIP\ (\nu^{-1})$ | Extrinsic (mosquito) incubation period | $1.03(4e^{5.15 - 0.123T})$ | (Caminade et al., 2017; Metelmann et al., 2021) |
| $r$ | Human recovery rate for dengue | $1/7$ | (Caminade et al., 2017) |
| $b$ | Probability of transmission of the disease from an infected vector to a susceptible host | $0.5$ | (Blagrove et al., 2020) |
| $B$ | Probability of transmission of the disease from an infected host to a susceptible vector | $0.31$ | (Metelmann et al., 2021) |

The definition of the weather and environmental variables is provided in Table SI2.

*Table SI2 – Weather and environmental variables and their definitions*

| **Weather and environmental variable** | **Meaning** |
|---|---|
| $T$ | Instantaneous temperature (°C), approximated using a sinusoidal function between on daily minimum and maximum (Metelmann et al., 2019) |
| $\bar{T}$ | Mean daily temperature (°C) |
| $T_7$ | Mean temperature of the last week (°C) |
| $T_w$ | Lowest winter (December-January-February) temperature (°C) |
| $R$ | Daily rainfall (mm) |
| $H$ | Human host density (hab/km$^2$); the original format of the GPWv4 data (Doxsey-Whitfield et al., 2015) is absolute count, but we converted this into density (people/km$^2$) to be consistent with the model input requirements |
| $P$ | Photoperiod (hours) |
| $L$ | Latitude (°) |

#### Exposure space

The methodology about the exposure space is depicted in Figure SI1.

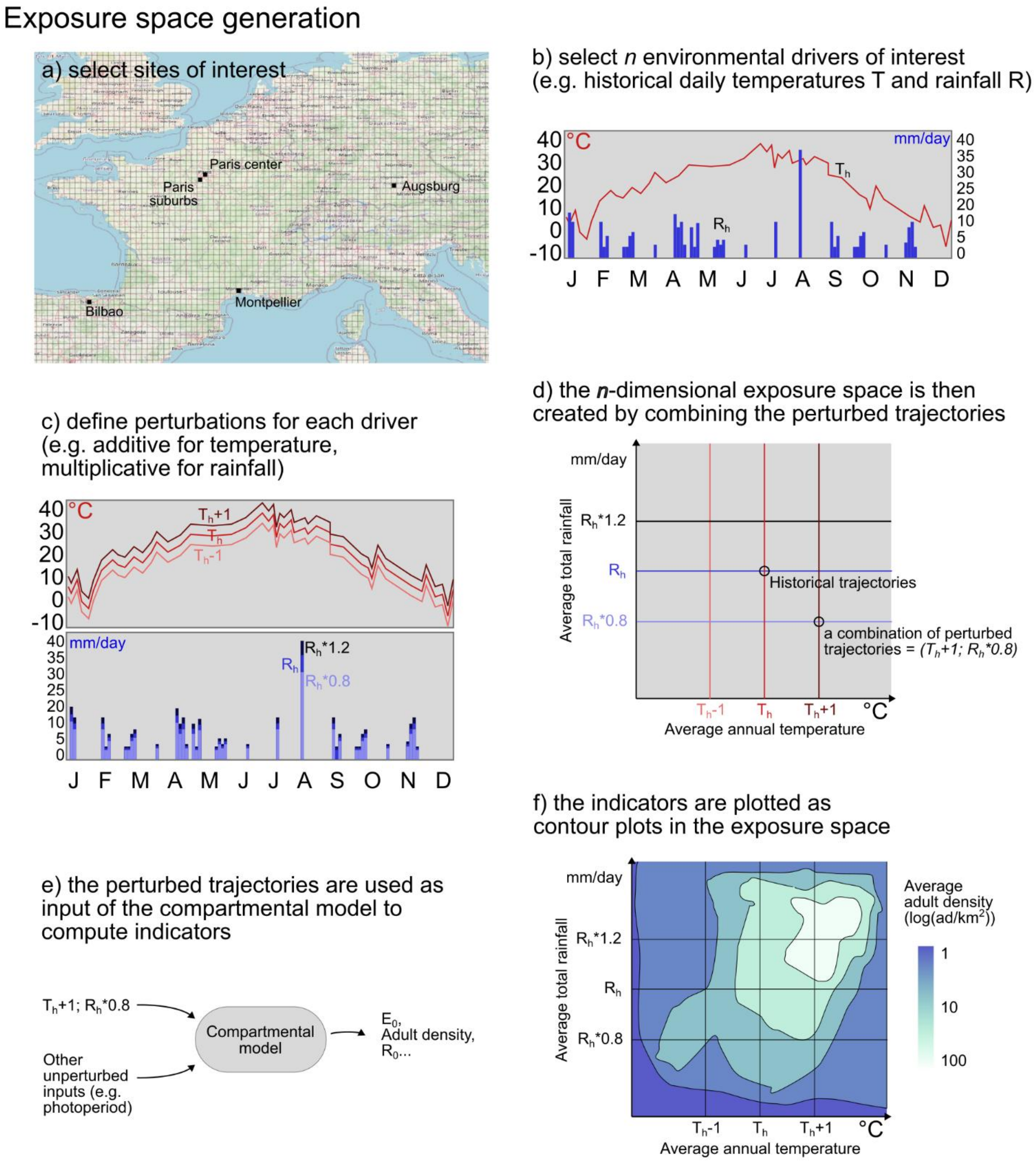


*Figure SI1 - Procedure to calculate the exposure space by perturbing trajectories of two environmental driving variables*.

### Additional results

#### Longitudinal ovitrap time series

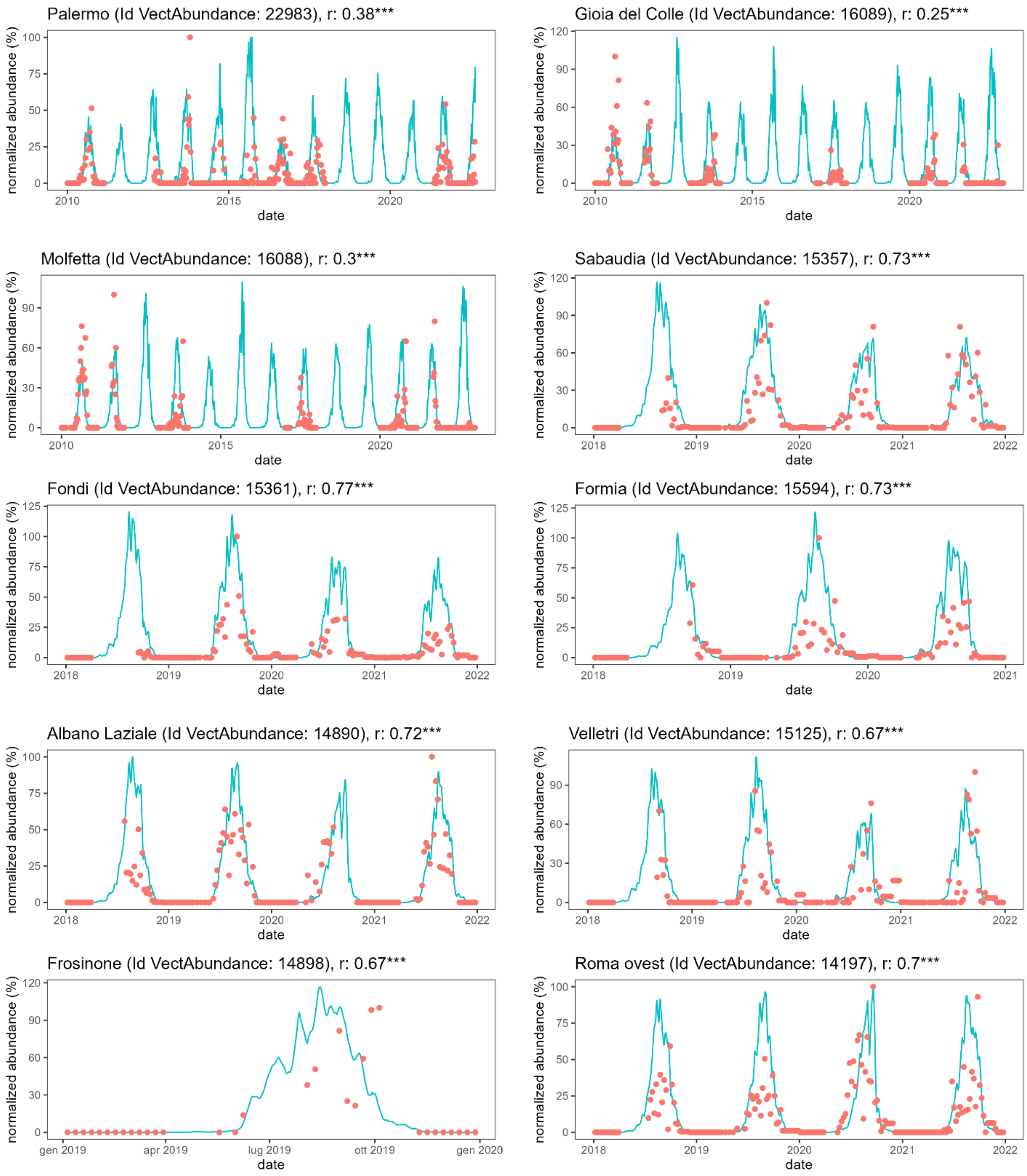

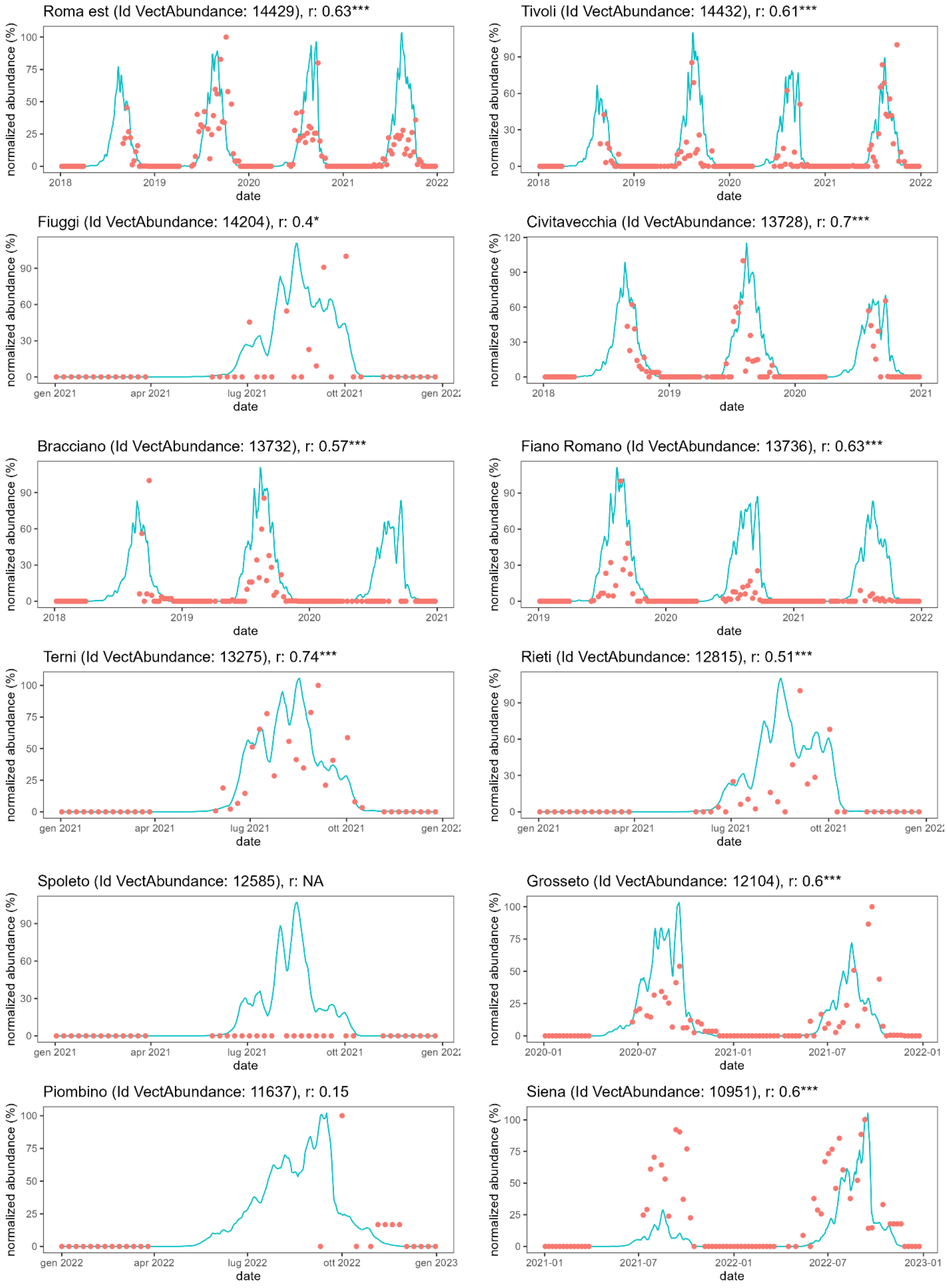
Roma est (Id VectAbundance: 14429), r: 0.63***
Tivoli (Id VectAbundance: 14432), r: 0.61***
Fiuggi (Id VectAbundance: 14204), r: 0.4*
Civitavecchia (Id VectAbundance: 13728), r: 0.7***
Bracciano (Id VectAbundance: 13732), r: 0.57***
Fiano Romano (Id VectAbundance: 13736), r: 0.63***
Terni (Id VectAbundance: 13275), r: 0.74***
Rieti (Id VectAbundance: 12815), r: 0.51***
Spoleto (Id VectAbundance: 12585), r: NA
Grosseto (Id VectAbundance: 12104), r: 0.6***
Piombino (Id VectAbundance: 11637), r: 0.15
Siena (Id VectAbundance: 10951), r: 0.6***
normalized abundance (%)
date
2018
2019
2020
2021
2022
gen 2021
apr 2021
lug 2021
ott 2021
gen 2022
apr 2022
lug 2022
ott 2022
gen 2023
2020-01
2020-07
2021-01
2021-07
2022-01
2022-07
2023-01

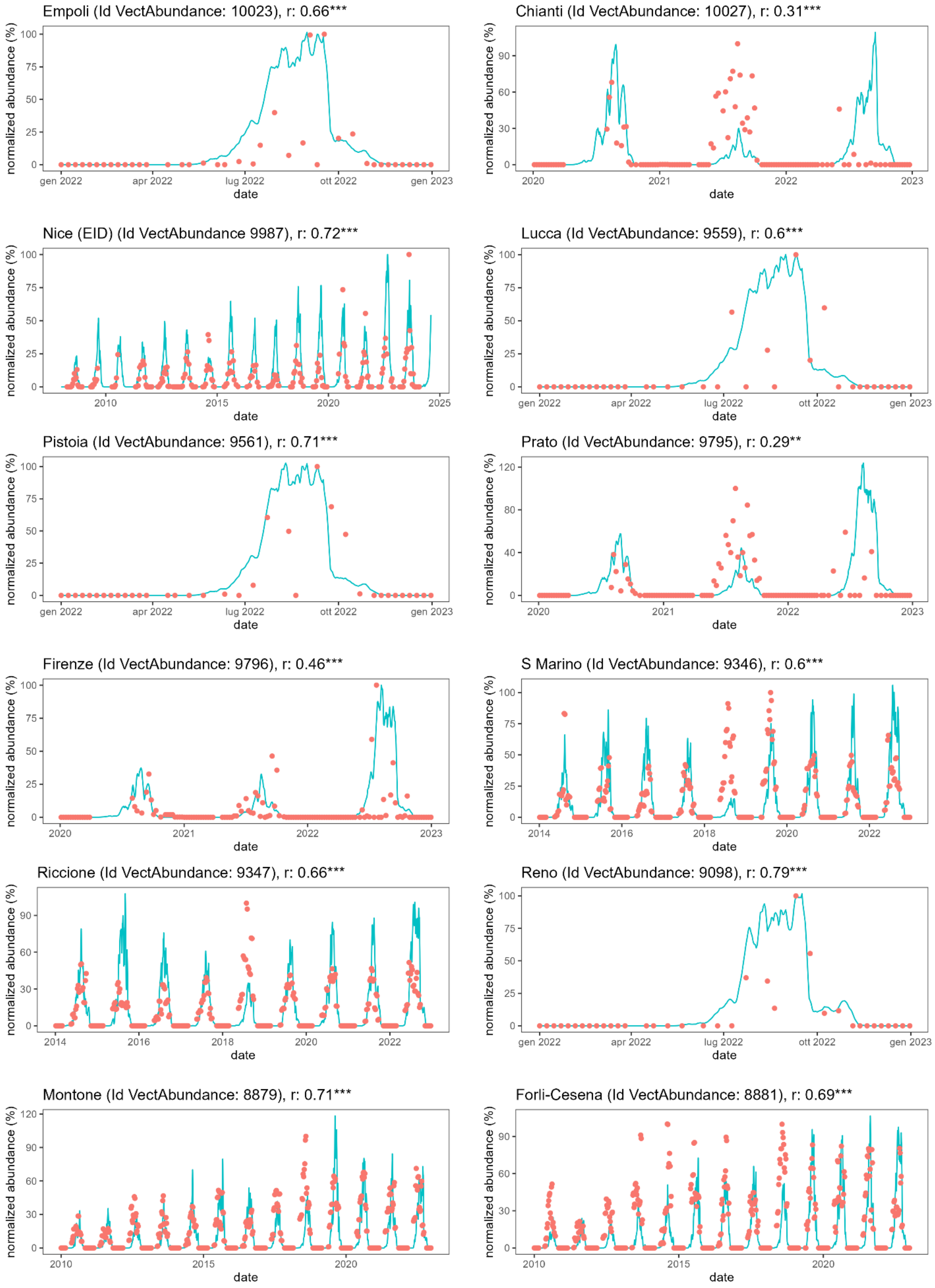
Empoli (Id VectAbundance: 10023), r: 0.66***
Chianti (Id VectAbundance: 10027), r: 0.31***
Nice (EID) (Id VectAbundance 9987), r: 0.72***
Lucca (Id VectAbundance: 9559), r: 0.6***
Pistoia (Id VectAbundance: 9561), r: 0.71***
Prato (Id VectAbundance: 9795), r: 0.29**
Firenze (Id VectAbundance: 9796), r: 0.46***
S Marino (Id VectAbundance: 9346), r: 0.6***
Riccione (Id VectAbundance: 9347), r: 0.66***
Reno (Id VectAbundance: 9098), r: 0.79***
Montone (Id VectAbundance: 8879), r: 0.71***
Forli-Cesena (Id VectAbundance: 8881), r: 0.69***
normalized abundance (%)
date
gen 2022
apr 2022
lug 2022
ott 2022
gen 2023
2010
2014
2015
2016
2018
2020
2021
2022
2023
2025

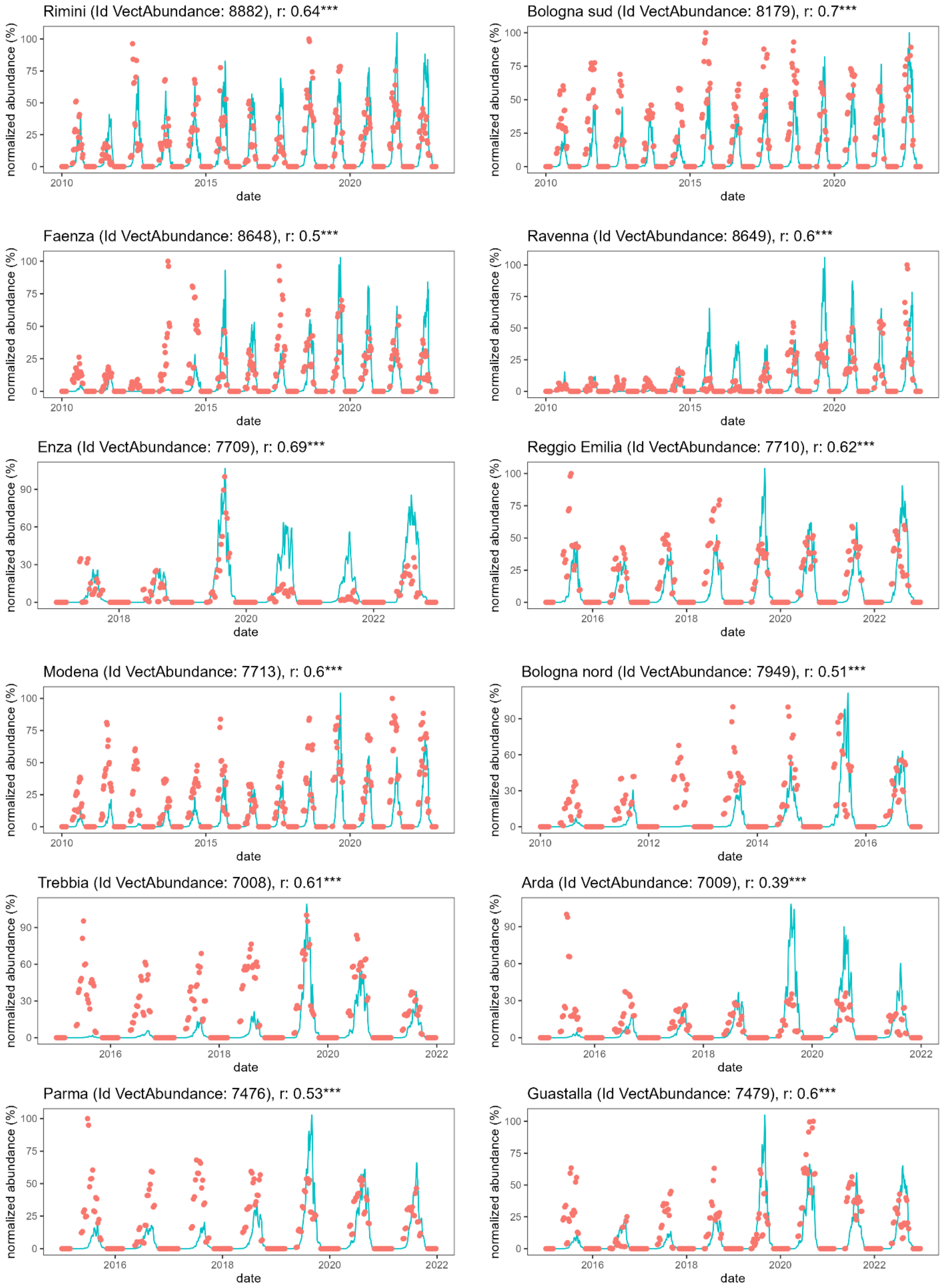

Rimini (Id VectAbundance: 8882), r: 0.64***
Bologna sud (Id VectAbundance: 8179), r: 0.7***
Faenza (Id VectAbundance: 8648), r: 0.5***
Ravenna (Id VectAbundance: 8649), r: 0.6***
Enza (Id VectAbundance: 7709), r: 0.69***
Reggio Emilia (Id VectAbundance: 7710), r: 0.62***
Modena (Id VectAbundance: 7713), r: 0.6***
Bologna nord (Id VectAbundance: 7949), r: 0.51***
Trebbia (Id VectAbundance: 7008), r: 0.61***
Arda (Id VectAbundance: 7009), r: 0.39***
Parma (Id VectAbundance: 7476), r: 0.53***
Guastalla (Id VectAbundance: 7479), r: 0.6***
normalized abundance (%)
date

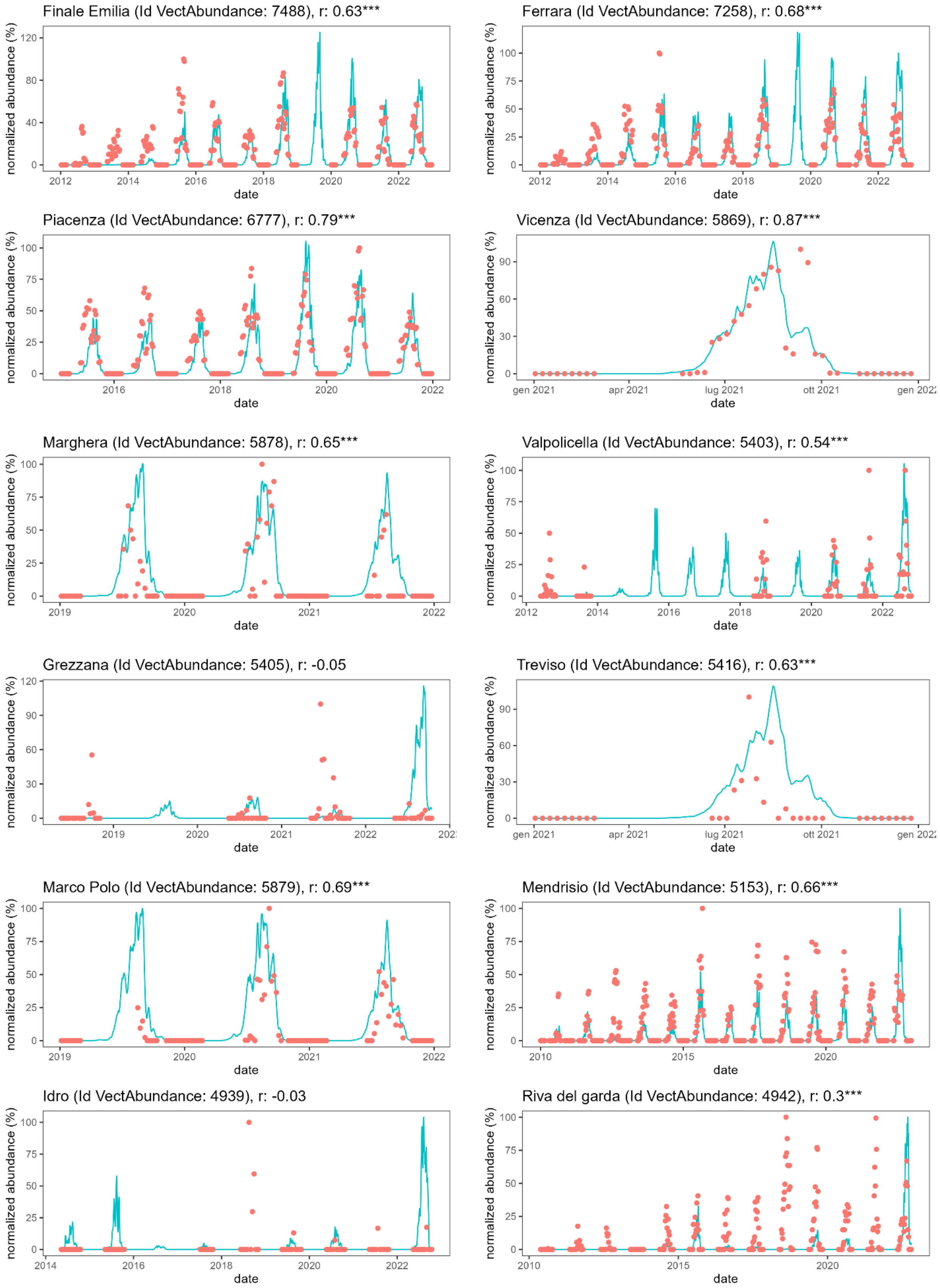

Finale Emilia (Id VectAbundance: 7488), r: 0.63***
Ferrara (Id VectAbundance: 7258), r: 0.68***
Piacenza (Id VectAbundance: 6777), r: 0.79***
Vicenza (Id VectAbundance: 5869), r: 0.87***
Marghera (Id VectAbundance: 5878), r: 0.65***
Valpolicella (Id VectAbundance: 5403), r: 0.54***
Grezzana (Id VectAbundance: 5405), r: -0.05
Treviso (Id VectAbundance: 5416), r: 0.63***
Marco Polo (Id VectAbundance: 5879), r: 0.69***
Mendrisio (Id VectAbundance: 5153), r: 0.66***
Idro (Id VectAbundance: 4939), r: -0.03
Riva del garda (Id VectAbundance: 4942), r: 0.3***
normalized abundance (%)
date
gen 2021
apr 2021
lug 2021
ott 2021

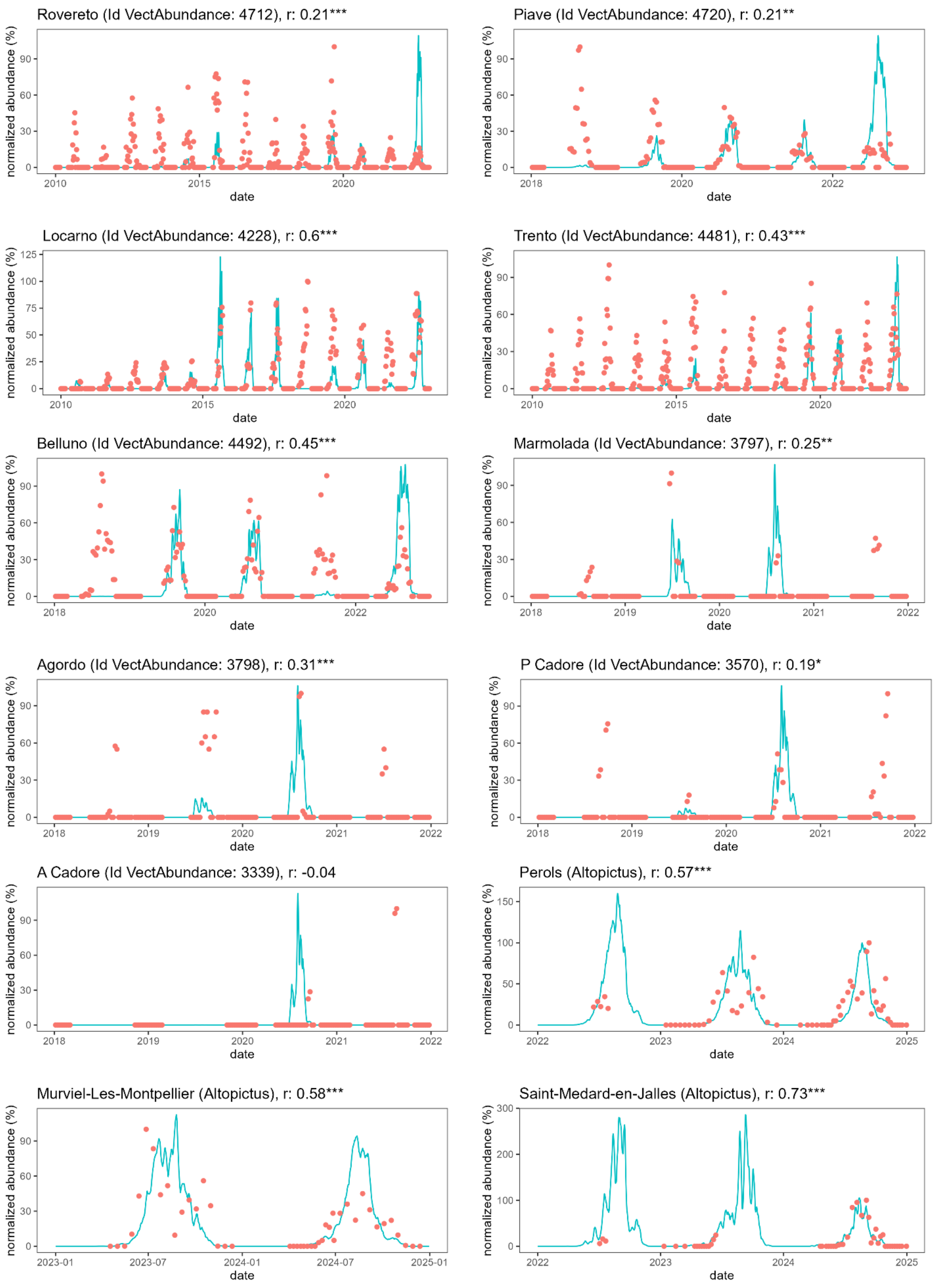
Rovereto (Id VectAbundance: 4712), r: 0.21***
Piave (Id VectAbundance: 4720), r: 0.21**
Locarno (Id VectAbundance: 4228), r: 0.6***
Trento (Id VectAbundance: 4481), r: 0.43***
Belluno (Id VectAbundance: 4492), r: 0.45***
Marmolada (Id VectAbundance: 3797), r: 0.25**
Agordo (Id VectAbundance: 3798), r: 0.31***
P Cadore (Id VectAbundance: 3570), r: 0.19*
A Cadore (Id VectAbundance: 3339), r: -0.04
Perols (Altopictus), r: 0.57***
Murviel-Les-Montpellier (Altopictus), r: 0.58***
Saint-Medard-en-Jalles (Altopictus), r: 0.73***
normalized abundance (%)
date

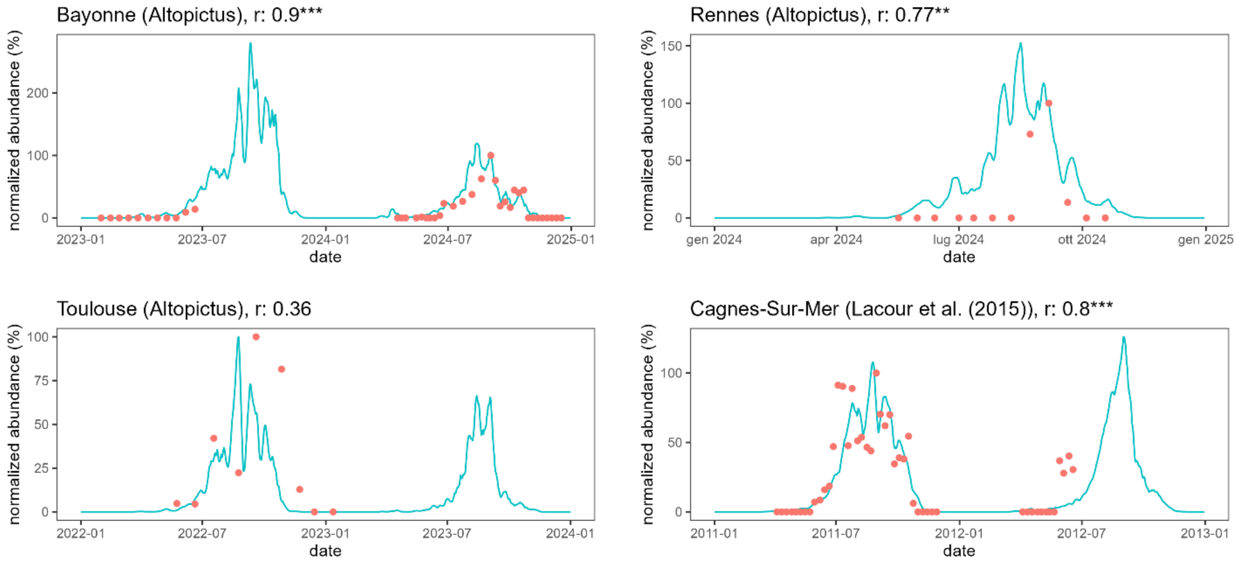


*Figure SI2 - Comparison between observed (orange dots) and simulated (blue line) standardized laid eggs rate per day (%) for different European locations. Correlations coefficients and associated statistical significance levels are provided in titles.*
